# Integration of cell-specific gene expression and chromatin accessibility facilitates localization of neurodegenerative risk in microglia

**DOI:** 10.64898/2026.09.15.751821

**Authors:** Xylena Reed, Dominic J. Acri, Alexandra Beilina, Jinhui Ding, Sara Saez-Atienzar, Mary Kaileh, Sarah Bromberek, Fangle Hu, Grigoriy Lerner, Daniel M. Ramos, Sultana Solaiman, Makayla Portley, Cory A. Weller, D. Thad Whitaker, Debra J. Ehrlich, Luigi Ferrucci, Sonja W. Scholz, J. Raphael Gibbs, Mark R. Cookson

## Abstract

Genome-wide association studies (GWAS) have identified many loci that contribute to the risk of neurodegenerative diseases. However, a persistent challenge in interpretation of GWAS is to break loci down to specific genes, variants, and cell types, and thus nominate disease mechanisms. Here, we used iPSC-derived cells containing population-level variation to examine GWAS loci across NDDs including Alzheimer’s disease, Parkinson’s disease and Lewy body dementia. We differentiated a set of 135 iPSC donor lines into two cell types relevant to neurodegeneration, neurons and microglia, and completed single cell gene expression and chromatin accessibility profiling. Meta-analysis of these data with published human brain snRNAseq for QTL mapping identified multiple loci associated with NDDs that are restricted to either neurons or microglia. Colocalization of GWAS and these QTL supports microglia as having a strong contribution to disease risk. We tested peaks nominated at the *BIN1* locus for enhancer activity using a perturb-seq-based method in microglia. Our results show one of the nominated peaks controls *BIN1* expression in microglia but also modifies expression of other genes at the locus. These results support the hypothesis that common variants affecting gene expression specifically in microglia can contribute directly to NDD risk rather than functioning solely as a secondary response to neurodegeneration. These data also show that iPSC-derived cells are a useful model to experimentally dissect GWAS loci that colocalize with QTL.

## Introduction

Genome-wide association studies (GWAS) have been successful in nominating genomic loci underlying risk of common phenotypes, including neurodegenerative diseases (NDDs). For example, Alzheimer’s disease (AD) risk GWAS have identified variants at 38 independent loci that are associated with disease (Wightman *et al*., 2021; Bellenguez *et al*., 2022). Similarly, meta-analyses of Parkinson’s Disease (PD) have identified 90 independent loci across the genome (Nalls *et al*., 2019) in European samples, with additional loci identified in Asian (Foo *et al*., 2020) and African populations (Rizig *et al*., 2023). Finally, Lewy Body Dementia (LBD), a disease with pathological features partially overlapping both AD and PD, has been shown to share genetic loci of both diseases (Chia *et al*., 2021; Scholz *et al*., 2025). Thus, large-scale genetic analyses have been successful in identifying the genetic risk of several interrelated neurological disorders. However, the interpretation of signals from GWAS is complex and often ambiguous. Due to linkage disequilibrium (LD), each nominated locus typically contains multiple genetic variants that could be functionally associated with multiple genes. This makes it difficult to resolve loci into more discrete units, such as variants, transcription factor binding sites or specific genes, using genetic information alone. Therefore, multiple strategies have been used to prioritize genes and variants within loci, largely using colocalization of disease risk loci with quantitative trait loci (QTL) maps derived from relevant tissue samples (Lappalainen *et al*., 2024). Commonly used approaches consider the relationship between genetic variants and RNA expression (eQTL) or epigenetic marks such as CpG methylation (methQTL) or chromatin accessibility (caQTL) (Ohlei et al., 2023; Xiong et al., 2023; Zeng et al., 2022). In principle, complex multimodal maps could be used to draw a mechanistic line between genetic variants and proximate biological events that would be predicted to contribute to disease. For heterogeneous tissues, including the brain, such QTL maps can be supplemented with cellular enrichment approaches to identify which cells are likely mediators of disease processes.

A limitation of these approaches is that they are correlational in nature and require additional experimental confirmation. There are multiple examples showing that iPSC models may be useful in interrogating endogenous human variation at specific loci (Soldner et al., 2016; Nott et al., 2019). Additional studies have used iPSC models to look at QTLs in specific cell types at a genome-wide scale (Schwartzentruber *et al*., 2018; Jerber *et al*., 2021; Bressan *et al*., 2023; D’Antonio *et al*., 2023). In addition, we have shown, using single-cell eQTL mapping in human brain, that a locus on chromosome 12 associated with PD risk is also correlated with higher expression of the known Parkinson’s disease-linked gene, *LRRK2* in microglia and not in other cell types that express the same gene (Langston et al., 2022). Importantly, we also confirmed the same association in induced pluripotent stem cell (iPSC) derived microglia from donors with the same genotypes. In this specific example, we were able to nominate microglia as a likely effector cell for this locus and make an important mechanistic distinction between gene expression and cell-type restricted gene quantitative traits.

Here, we have generated a map of caQTL and eQTL in iPSC-derived cell types that are relevant in the development of neurodegenerative diseases, namely cortical neurons and microglia. Additionally, we show that iPSC-derived cell samples can be used to increase power when combined with existing single cell human brain data via meta-analyses. By adding chromatin accessibility and expression in a cell-specific manner, we were able to show that colocalization between GWAS signals and QTL. Finally, we use a risk-peak-directed modified perturb-seq protocol (Dixit *et al*., 2016) to functionally test the regulatory potential of five nominated peaks of gene expression at the AD- and LBD-associated locus encoding *BIN1*.

## Results

### Generation of single-cell expression and chromatin accessibility for iNeurons and iMicroglia

To generate neurons and microglia, we used a recently described cohort of iPSCs derived from healthy donors from the Genetic and Epigenetic Signatures of Translational Aging Laboratory Testing (GESTALT; *n*=89) study (Reed *et al*., 2024) supplemented with donors from the Baltimore Longitudinal Study of Aging (BLSA; *n*=8) (Ershler *et al*., 2005) and samples collected from the NIH Movement Disorders Research Clinic (*n*=41)(Table S1). The combined sample included 138 individual donors from multiple ancestry groups, with 56% being male (Figure S1). We genotyped each iPSC line to confirm genetic fidelity to its original donor, expanded lines individually to assess growth rates, then pooled lines into sets of six donors to limit the effects of differentiation batches on downstream analyses. Twenty-three pools were differentiated separately into forebrain neurons (iFBn) and microglia (iMGL) using previously published protocols (Burkhardt *et al*., 2013; Brownjohn *et al*., 2018; Reed *et al*., 2021) (Figure 1A).

**Figure 1.**
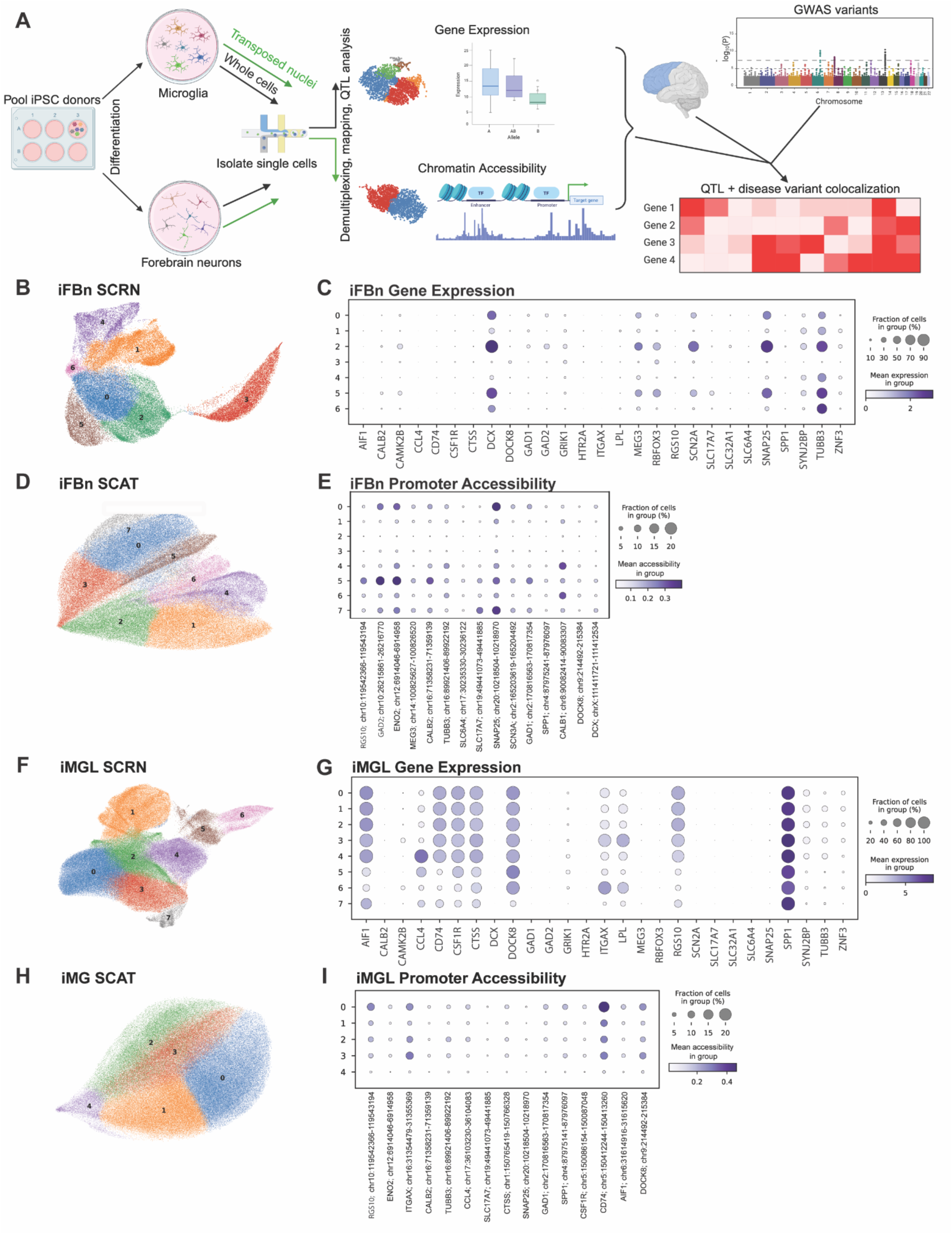
Experimental workflow and single cell validation of differentiation protocols. A) Study design from iPSC differentiation to microglia (iMGL) and forebrain neurons (iFBn) followed by single cell profiling of gene expression and chromatin accessibility, meta-analysis with data from human brain and colocalization with Genome wide association studies to nominate loci of interest. B) UMAP with Leiden based clustering at resolution 0.3 of scRNA-seq from iFBn. C) Mean expression of canonical neuronal and microglia markers for each cluster of iFBn. Size of circle indicates the fraction of cells expressing and color intensity indicates expression level per cell. D) UMAP of snATAC-seq from iFBn. E) Mean chromatin accessibility for regions around canonical gene markers for neurons in each iFBn cluster. Size of dot indicates fraction of cells with accessibility, and intensity indicates level of accessibility per cell. F) UMAP of scRNA-seq from iMGL. G) Dotplot showing expression of canonical neuronal and microglia markers in each cluster of iMGL. H) UMAP of snATAC-seq from iMGL. I) Dotplot showing chromatin accessibility for regions around canonical gene markers for microglia in each iMGL cluster.

After differentiation (day 60 for iFBn, or day 30 for iMGL), single-cell libraries were generated independently for both gene expression and chromatin accessibility in each cell type. Prior to clustering, single cell gene expression (SCRN) and single nuclei chromatin accessibility (SCAT) datasets were demultiplexed using genetic variants present in the individual donors. Genotype-based demultiplexing of these pools resulted in ∼80% of cells being assigned to a specific donor, with an average of approximately 13% of cells being ambiguous for donor and 7% assigned as doublets in the iFBn datasets. In the iMGL datasets, approximately 5% of cells were ambiguous for donor and 14% were called as doublets (Table S2). Three donors did not have a sufficient number of cells (>10) to be included in downstream analysis, resulting in 135 donors used for QTL mapping.

Gene expression and chromatin accessibility assays for each cell type were independently visualized on UMAPs, clustered using the Leiden algorithm and examined for canonical cell type markers (Figure 1 B-I). One gene expression cluster of iFBn (cluster 3) lacked the canonical neuronal markers *TUBB3* and *SNAP25*, and was excluded from further analyses due to assumed immaturity of the cells (Figure 1 B,C). In the SCAT dataset, iFBn clusters with low chromatin accessibility around the *SNAP25* promoter region were excluded (Figure 1 D,E). All iMGL clusters expressed *AIF1* (encoding the protein IBA1) and were therefore retained (Figure 1 F,G). However, iMGL SCAT showed a small subcluster (cluster 4) with low accessibility around the *AIF1* promoter in iMGL that was removed from downstream analyses (Figure 1H,I).

### Differentiated iFBn and iMGL expression profiles are similar to human brain cell types

Prior to performing QTL mapping and meta-analysis, we examined the correlation of overall gene expression between iFBn, iMGL and post-mortem cells from 248 donors in the publicly available Religious Orders Study and Memory and Aging Project (ROSMAP) post-mortem Dorso-Lateral Pre-frontal Cortex (DLPFC) data (Fujita *et al*., 2024). These cells were categorized into eight brain cell classes: Astrocytes (AST), Endothelial Cells (END), Excitatory neurons (ExN), Inhibitory Neurons (InN), Microglia (MGL), Oligodendrocytes (OLG), Oligodendrocyte Precursor Cells (OPC), and Pericytes (PER). The expression profiles of iFBn were most highly correlated with post-mortem DLPFC neuronal cell types (ExN and InN), whereas iMGL expression profiles were most similar to post-mortem DLPFC Microglia and DLPFC CNS Associated Macrophages (CAMs) (Figure 2A). Using Principal Component Analysis (PCA) of gene expression pseudobulked at the donor level the iMGL donors group closely with DLPFC Microglia (Figure 2B). iFBn were overlapped with DLPFC ExN on the same PCA plots but also occupied a broader transcriptional space, possibly indicating a relative lack of specialization in these cells compared to the complexity of mature human brain neurons. These comparisons suggest that iMGL are transcriptionally similar to human brain cells at a broad cell type level as shown previously (Brownjohn *et al*., 2018; Langston *et al*., 2022).

**Figure 2.**
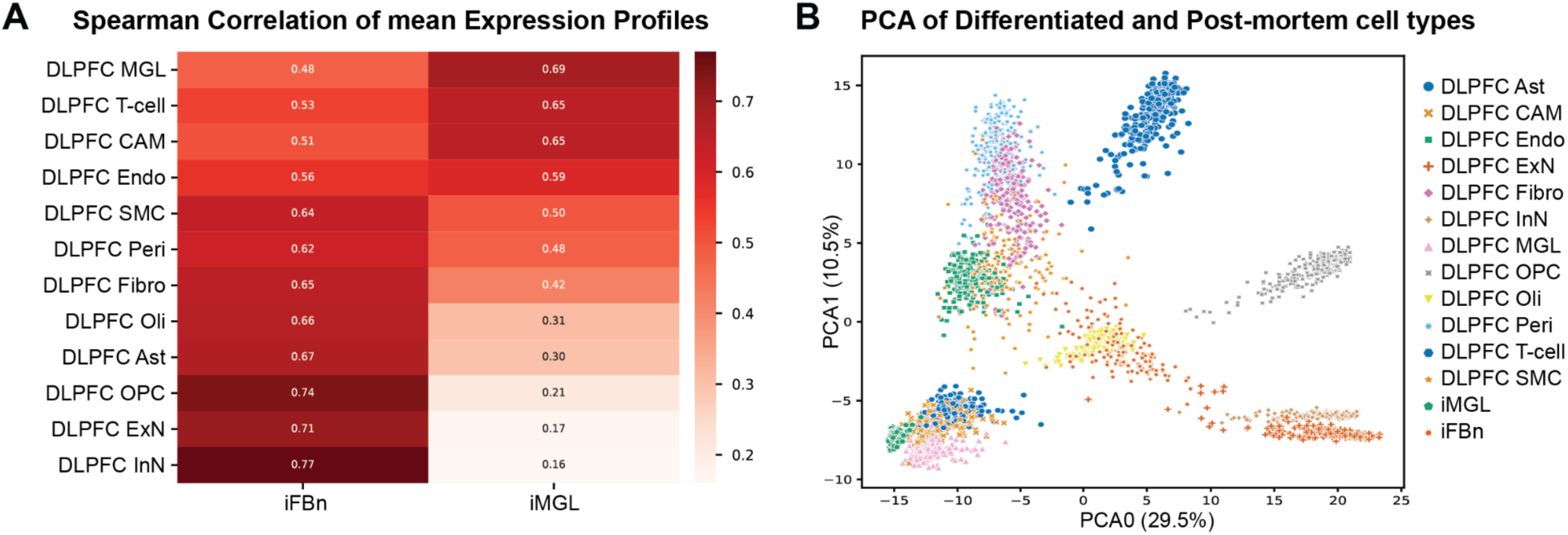
Correlation of iFBn and iMGL results with ROSMAP dataset. A) Spearman correlation of mean expression patterns between iFBn and iMGL and cell types from public datasets. B) PCA showing similarity between differentiated and post-mortem cell types. Each point represents one sample and the color and shape correspond to cell type.

### Identification of Quantitative Trait Loci (QTL)

Next, we generated QTL maps for each differentiated cell type to identify *cis*-QTL within +/- 1 Mb from the feature (peak for caQTL, gene for eQTL). We did not detect any genome-wide significant caQTL in iFBn but did identify 2,488 ATAC peaks with a nominally significant empirical p-value <0.05. In contrast, there were 2,125 caQTL detected in iMGL with 7580 nominally significant ATAC peaks. A similar difference in detection of QTL for expression was noted between cell types as we identified 27 eQTL in iFBn and 2,211 eQTL in iMGL (Table 1). There were only 11 eQTL that were significant in both cell types and no caQTL were shared between cell types. Using the same methods we identified eQTL in the ROSMAP DLPFC ExN, InN and MGL.

**Table 1.**
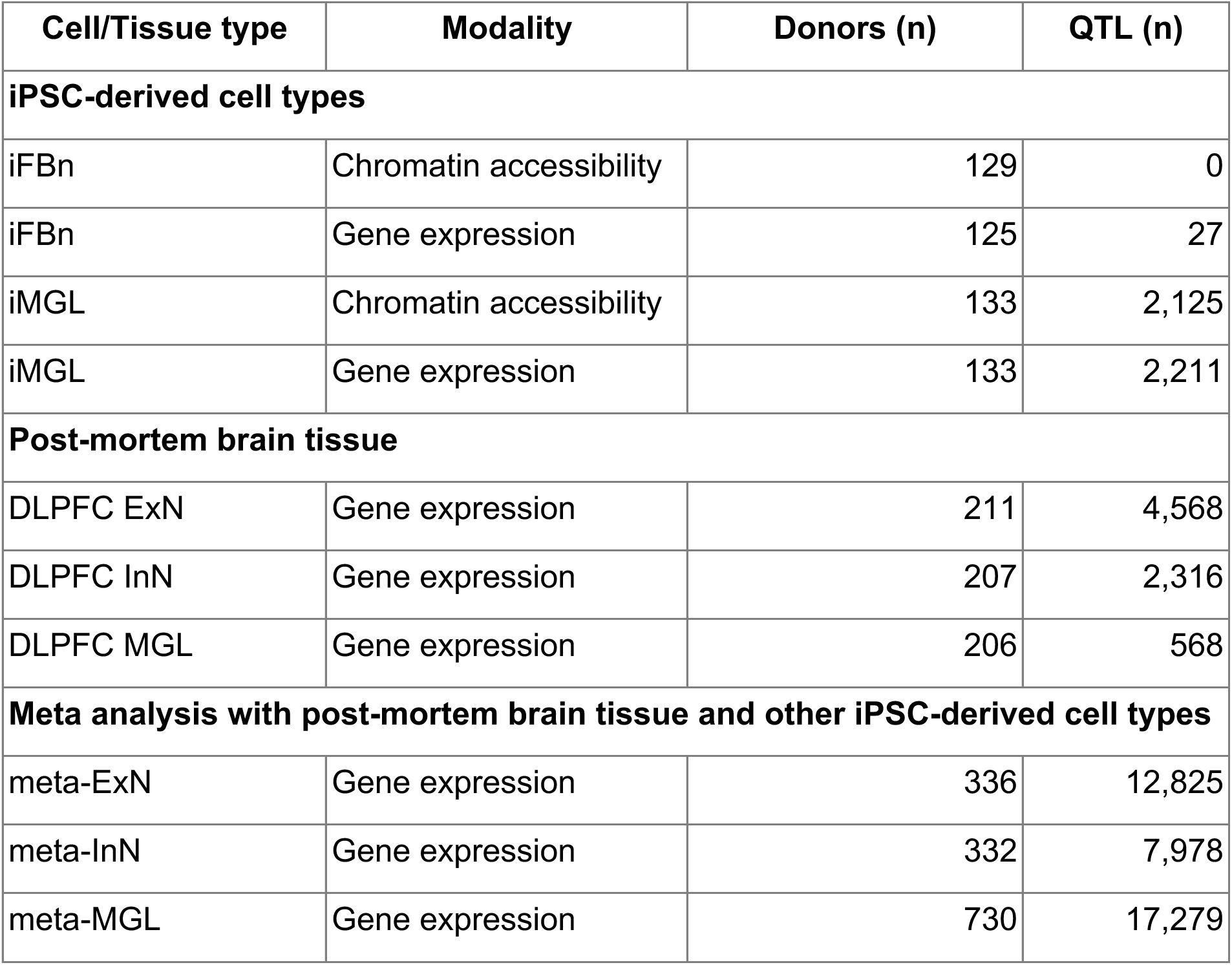
Number of QTL identified in each modality and cell type. iFBn and iMGL refer to new datasets generated as part of this study. Meta refers to the meta-analysis with all selected public datasets, and post-mortem refers to meta-analysis of post-mortem Dorso-Lateral Prefrontal Cortex (DLPFC) samples.

| Cell/Tissue type | Modality | Donors (n) | QTL (n) |
| --- | --- | --- | --- |
| **iPSC-derived cell types** |  |  |  |
| iFBn | Chromatin accessibility | 129 | 0 |
| iFBn | Gene expression | 125 | 27 |
| iMGL | Chromatin accessibility | 133 | 2,125 |
| iMGL | Gene expression | 133 | 2,211 |
| **Post-mortem brain tissue** |  |  |  |
| DLPFC ExN | Gene expression | 211 | 4,568 |
| DLPFC InN | Gene expression | 207 | 2,316 |
| DLPFC MGL | Gene expression | 206 | 568 |
| **Meta analysis with post-mortem brain tissue and other iPSC-derived cell types** |  |  |  |
| meta-ExN | Gene expression | 336 | 12,825 |
| meta-InN | Gene expression | 332 | 7,978 |
| meta-MGL | Gene expression | 730 | 17,279 |

*.Prioritization of genes using QTL colocalization with neurodegenerative diseases*

Results from both iFBN and iMGL were taken forward to meta-analyses to evaluate the hypothesis that addition of iPSC-derived data to data from human brain QTL maps would improve statistical power to detect QTLs. The same method of *cis*-eQTL analysis was performed on each of the eight cell types in the ROSMAP DLPFC dataset limiting to donors including only the donors with no cognitive impairment or with mild cognitive impairment (MCI) to minimize any potential effects of advanced neurological disease or no other sources of impairment (*n*∼200). To determine that this is a valid approach, prior to performing *cis*-eQTL meta-analysis, we performed a direct intersectional analysis of our eQTL results with public datasets. We identified eQTL in each study, and examined the overlap of the eQTL with our datasets (Table 1). Spearman correlation of iMGL was highest with post-mortem MGL at 0.49, and iFBn Spearman correlation was highest with post-mortem ExN and post-mortem InN at 0.39 and 0.38, respectively (Figure 3A). These analyses also reinforce the concept that iMGL were an appropriate model for eQTL mapping.

**Figure 3.**
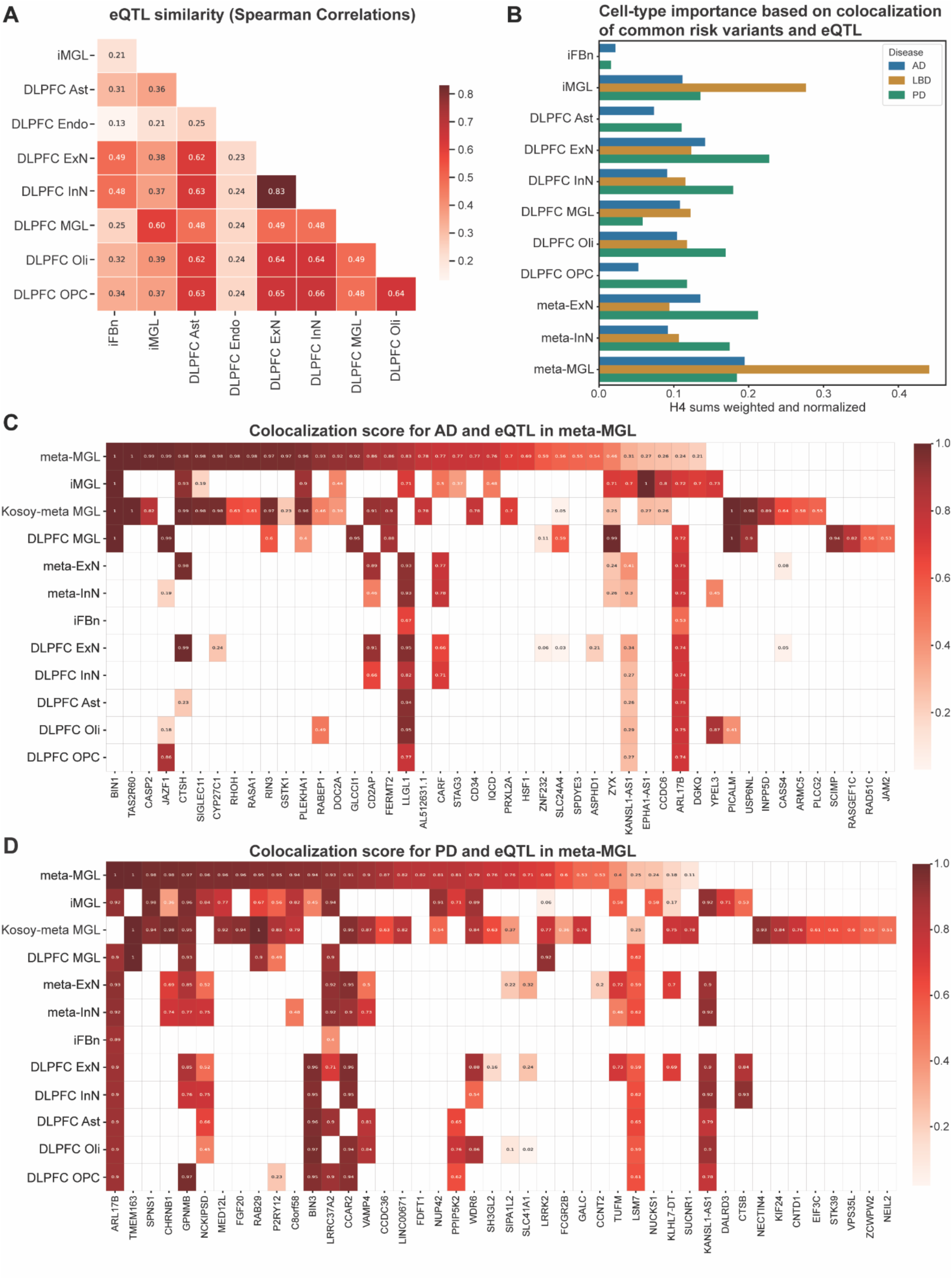
Functional prioritization of eQTL in MGL. A) Spearman correlation between eQTL identified in each cell type. Darker shade of red indicates higher correlation. B) Cell type importance for each NDD based on colocalization scores. Blue is AD, Orange is LBD and Green is PD. The x-axis shows the normalized and weighted H4 sums. C) AD risk genes from GWAS colored by colocalization with eQTL H4 score (darker red is more significant). D) Heatmap of PD risk genes from GWAS colored by colocalization H4 score (darker red is more significant). Colocalization results are filtered for protein coding genes that were detected in at least one MGL dataset. Full results are in Table S3.

### Prioritization of genes using QTL colocalization with neurodegenerative diseases

Next, we evaluated the utility of meta-QTL nominate mechanisms for neurodegenerative diseases (NDDs) based on GWAS summary statistics from three previously published studies for PD, LBD, and AD (Nalls *et al*., 2019; Chia *et al*., 2021; Bellenguez *et al*., 2022) using colocalization to identify genetic loci shared between each disease and iPSC-derived cell type eQTL (Giambartolomei *et al*., 2014). Specifically, we estimated posterior probability of the same genetic locus being associated with and shared between two different traits, in this case, eQTL and disease risk, reported as H4. This analysis identified loci in both cell types with support of shared genetic architecture between GWAS risk and QTL (H4 > 0.5) in each NDD diagnosis (Table 2; Table S3). Expanding to include meta-analysis with post-mortem brain datasets significantly increased the number of detected overlap for neuronal cell types, and increased the number of loci with colocalization support for each NDD in meta-MGL by 2-3 fold.

**Table 2.**
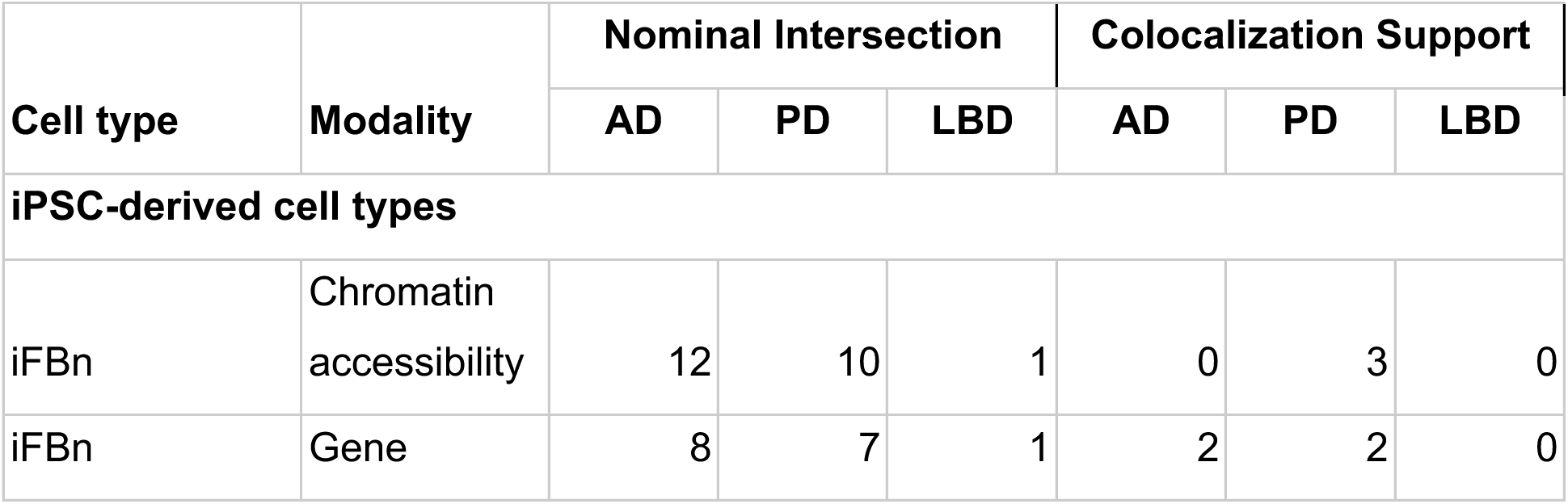
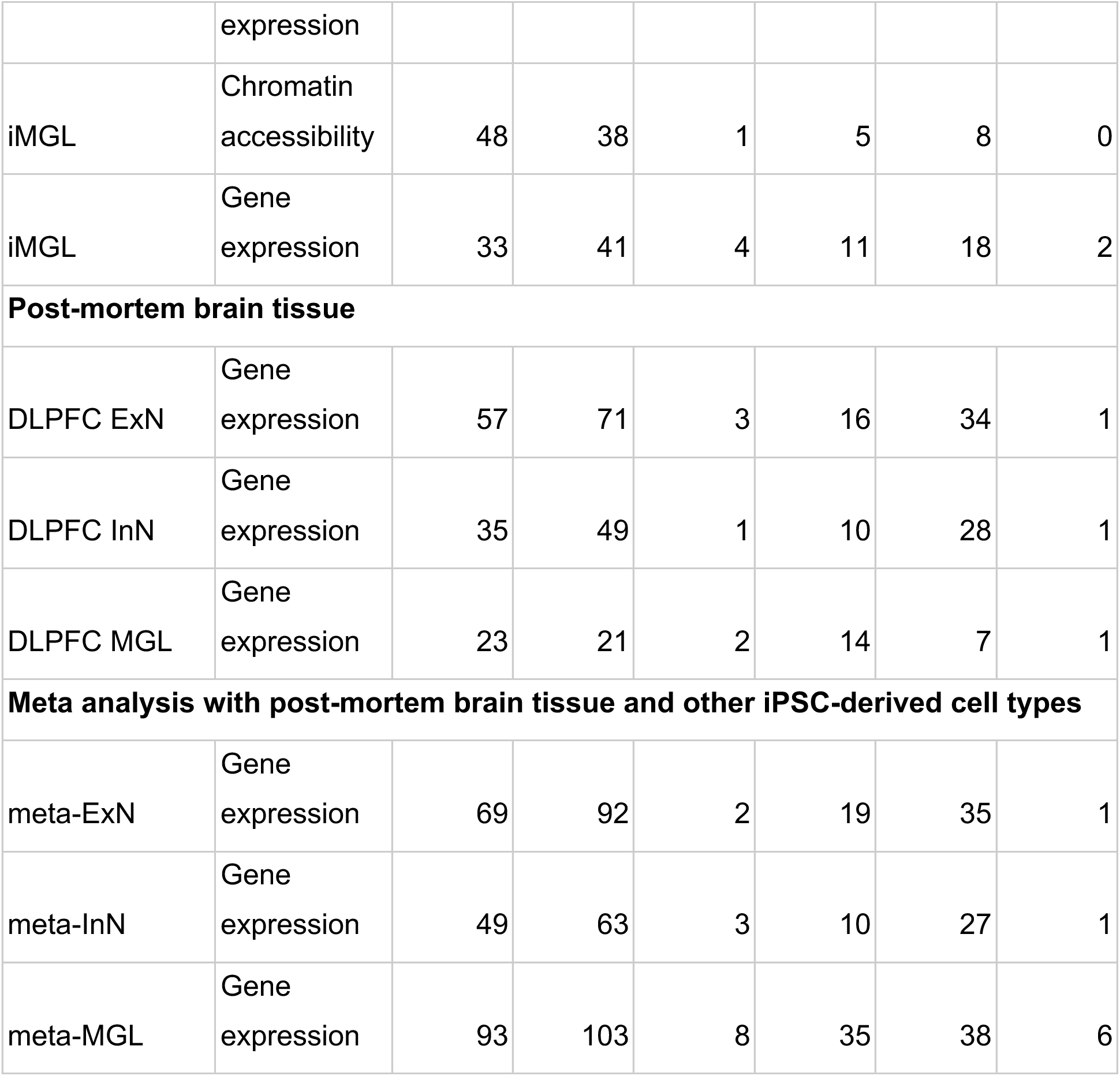
Number of QTL from each modality and cell type having nominal intersection with GWAS variants or Colocalization support.

Examination of cell-type importance for each NDD demonstrated that multiple cell types including neurons and microglia contribute to risk of disease (Figure 3B). This is notably in contrast to gene expression alone, which has nominated microglia for AD (Bellenguez *et al*., 2022) and neurons (Kamath *et al*., 2022) or oligodendrocyte precursor cells (Agarwal *et al*., 2020) for PD.

AD risk loci intersected with iMGL eQTL for 33 genes and had colocalization support at 11 genes. The number of intersections was 23 and 93 genes, for post-mortem MGL and meta-MGL respectively, and the number of genes with colocalization support also increased to 14 (post-mortem MGL) and 35 (meta-MGL), indicating that additional samples increase the power to identify relevant eQTL (Table 2). The gene with the highest colocalization score between AD and MGL across all analyses was *BIN1* on chromosome 2 (Figure 3C). The *BIN1* colocalization was strong and robust in MGL for both the meta-eQTL and all included substudies (H4 from 0.999 to 1), and was not identified in any other analyzed cell type. Variants at the *BIN1* locus were first associated with AD in 2010 by GWAS (Seshadri *et al*., 2010) and a signal was also identified in a recent genome sequencing study of patients with LBD (Chia *et al*., 2021). Previous studies have also reported this strong colocalization between AD risk and microglia eQTL for BIN1 (Kosoy *et al*., 2022; Lopes *et al*., 2022). PLEKHA1 also showed significant colocalization across MGL studies (H4=0.4 - 0.96; Figure 3C and Figure S2) and has previously been reported to have higher expression in immune cells from AD post-mortem brain compared to controls (Lindbohm *et al*., 2026). There were eight iFBn eQTL genes that intersected with AD GWAS, and two with colocalization support (Table S3). However, the number of intersections in the post-mortem and meta-analysis of both ExN and InN were much higher than in iFBn alone (Table 2). The strongest signals that were specific to neuronal cell types were *PRSS36* (H4>0.98; Figure S3A), and *ACE* (H4>0.94; Figure S3B) where eQTL signal was higher in ExN for *PRSS36* and higher in InN for *ACE* (Table S3). Interestingly, both loci contained iMGL specific risk peaks, however the eQTL signal was much lower in MGL than neurons.

In PD, we identified 41 nominal intersections for iMGL, and 18 that had colocalization support (Table S3). Consistent with our previous data (Langston *et al*., 2022), the PD gene *LRRK2* showed an eQTL across all microglia results and it displayed strong colocalization between risk and the meta-microglia and post-mortem microglia results, however, the colocalization with iMGL was poor (Figure 3D, Figure S4). Additionally *RAB29* showed strong and consistent microglia-specific colocalization with PD risk (Figure 3D and Figure S5) and has previously been functionally linked with *LRRK2* (MacLeod et al., 2013; Beilina et al., 2014; Purlyte et al., 2018). Colocalization with risk at an eQTL for *GPNMB*, which was recently suggested to bind to ɑ-synuclein and identified as a marker of lysosome dysfunction (Diaz-Ortiz et al., 2022; Bogacki et al., 2025), was strong across analyses and present in multiple cell types including microglia, neurons and oligodendrocyte precursors (Figure 3D). There was strong MGL-specific colocalization support at *SPNS1*, which has not previously been nominated as a risk gene, that highlighted a potential coding variant in nearby gene CD19 (CADD = 10.71), (Figure 3D and Figure S6). *P2RY12*, previously nominated as a risk gene for PD (Andersen *et al*., 2021; Lopes *et al*., 2022), also showed a strong eQTL risk colocalization in meta-microglia (H4 = 0.95), mixed in post-mortem MGL datasets (H4 = 0.85 and 0.49), but only weak support in iMGL (H4 = 0.56), as well as some signal in OPCs (H4=0.23) (Figure 3D).

We identified four intersections in our iMGL data with common risk in LBD, with two showing colocalization support (Figure S7; Table S3). Post-mortem MGL found *BIN1* as having colocalization support, while the meta-MGL analysis showed six loci with colocalization support, including two uncharacterized genes near *BIN1* (*RNU6-675P* and *AC012508.1*). Of these intersections, *BIN1* was the only gene that showed colocalization (H4 = 1.0) across all MGL datasets and appeared to be MGL specific. In neurons, *FGFRL1* was colocalized with risk in ExN and InN but was not detected in the iFBn dataset.

Using the same method, we examined caQTL colocalization with disease risk in iMGL and iFBn. From this analysis we identified 48 loci in AD iMGL that showed nominal intersection with caQTL, where five had colocalization support (Table S4). Of these peaks, one was found in an intron of *PLEKHA1*, which we showed above also had an eQTL with colocalization support with AD (Supp. Fig 2A-B). A second colocalized peak was found upstream of *PICALM*, where we saw eQTL colocalization support in the two DLPFC MGL datasets, but not in our iMGL data (Figure 3C). iFBn caQTL showed intersection at 12 loci, but none had colocalization support in AD. PD risk intersected with iMGL caQTL at 38 ATAC peaks and showed colocalization support with PD risk at eight peaks. Two of the colocalized peaks were located near the *TMEM163* locus, where we also saw an eQTL colocalized with PD in all MGL analyses except iMGL (Figure 3D). iFBn caQTL intersected with PD risk at 10 ATAC peaks with support for colocalization with risk at three peaks. Two of these overlapped two alternate transcription start sites of *KANSL1*, and one also covered the predicted start site of *KANSL1-AS1* which we found to be colocalized with eQTL and PD-risk across multiple cell types (Figure 3D). iMGL caQTL intersected with LBD risk at one ATAC peak near the *BIN1* locus; however, no support for colocalization with risk was detected. One iFBn caQTL also intersected with shared AD and LBD risk at a different ATAC peak near *SDSL*, where eQTL intersection had also been observed in our data (Table S3), but no support for colocalization with risk was detected for either the eQTL or the caQTL.

### Correlations between cis-peaks and genes

We next sought to establish whether our chromatin accessibility data can be used to further nominate potentially causative variants from neurodegenerative disease GWAS. To this end, we first examined the relationship between chromatin accessibility and gene expression in differentiated iPSCs by analyzing the correlation between the caQTL and eQTL within each cell type. We identified 5,136 genes in iFBn and 11,096 genes in iMGL that showed a statistically significant correlation with one or more *cis-*proximal peaks (+/- 1 Mb of a gene transcription start site, TSS). Of these, 3,105 genes were present in the results for both iFBn and iMGL, and 7,800 peak-gene pairs. In iFBn, 22.2% (6 of 27) of eQTL genes had a correlated cis-proximal ATAC peak, while 61.0% (1,349 of 2,211) of iMGL eQTL genes had a correlated *cis-*proximal peak, and 37.0% (787 of 2,125) of caQTL peaks were correlated with a *cis*-proximal gene.

Restricting the *cis-*correlated pairs to only those where the gene and the ATAC peak both have a detected QTL, the percentage of iMGL eQTL genes with *cis-*proximal intersection is 14.0% (309 of 2211) and, similarly, the iMGL caQTL intersection is 14.5% (308 of 2,125) with eQTL genes. However, based on a colocalization analysis between eQTL and *cis*-proximal caQTL in iMGL, only 12.4% (274 genes) of these possible pairings have support (posterior probability that locus is shared, H4 > 0.5) for colocalization of the QTL. This result suggests that while cell-type specific peaks may account for cell-specific *cis*-eQTL, only a small portion of these are driven by caQTL at these cell-type specific peaks.

We then filtered the correlated *cis*-proximal results for peaks containing variants that are associated with AD, PD and LBD. There were 218 iMGL peaks containing AD associated variants, correlated with 429 *cis*-proximal genes and 68 iFBn peaks associated with 91 AD genes (Table S5). This included two peaks where *BIN1* was identified as a *cis*-proximal gene, one in iFBn and one in iMGL. Based on PD risk variants, there were 180 iMGL peaks associated with 382 genes, and 74 iFBn peaks associated with 91 genes. For LBD GWAS risk variants, there were 11 iMGL peaks associated with 40 genes and two iFBn peaks associated with two genes in LBD. For a limited set of loci, we performed mediation analysis to determine if the eQTL signal was mediated by a caQTL signal. For seven loci (two AD and five PD) where disease risk is colocalized with eQTL in iMGL and is consistent with all studies used in our analyses we detected no mediation of an eQTL by a caQTL. These results suggest that regulation of gene expression is complex, and not due to simple alignment of caQTL and eQTL.

### Nomination of regions near BIN1 for functional validation

Since most GWAS loci are found in non-coding regions, we postulated that the candidate regions identified here may affect disease pathogenesis through modulation of *cis-*regulatory elements. To investigate this possibility, we took the genes with colocalization support above (Table 2, Table S3) and filtered our gene list to include targets that had evidence of association in more than one GWAS and had peaks that did not intersect with the proximal promoter for that gene. This analysis nominated the *BIN1* locus, which is a risk factor for both AD and LBD as discussed above, as having multiple potential *cis-*regulatory elements relevant for control of gene expression. We computed the colocalization score for genes at this locus across the MGL datasets and found that *BIN1* had an H4 = 1 across all MGL datasets, but no colocalization in other cell types, while other genes at the locus showed colocalization in some but not all of the MGL analyses (Figure 4A). The top variant in the region, rs6733839, showed significant effects as an eQTL for *BIN1* expression in all MGL included in our analysis (Figure 4B). Additionally the minor allele was associated with higher *BIN1* expression iMGL samples generated in this study (Figure S8A) and in the human brain, specifically the DLPFC from the ROSMAP study (Figure S8B).

**Figure 4.**
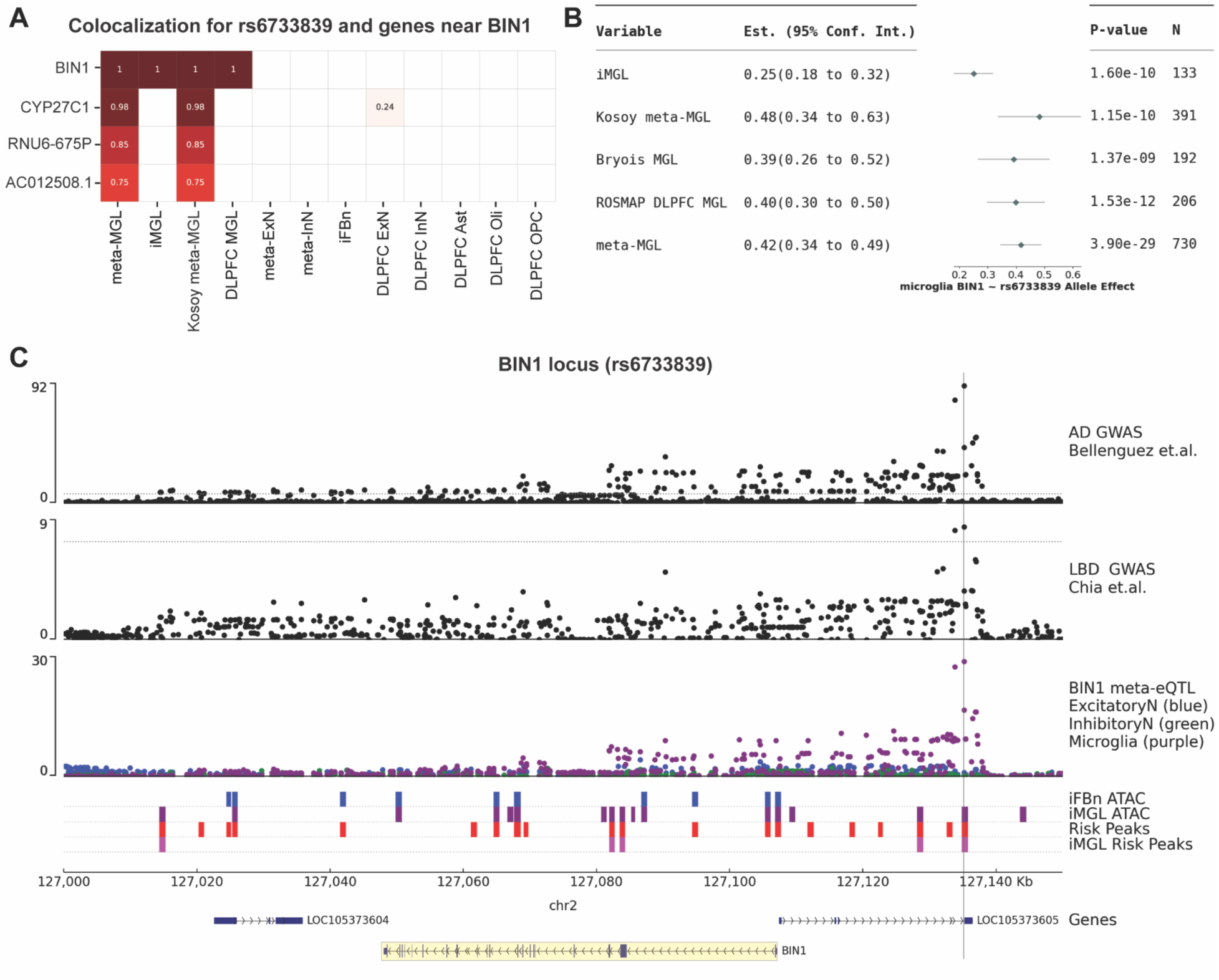
Nomination of peaks near BIN1 for functional analysis *in vitro*. A) Colocalization plot for AD and LBD risk near BIN1. B) Forest plot showing the effect of the lead SNP rs6733839, at the *BIN1* locus in each MGL analysis. C) Locus plot showing AD (top) and LBD (bottom) GWAS signal across the *BIN1* locus, with *BIN1* meta-eQTL signal below (ExN = blue, OnN = green, MGL = purple). Bars show the location of iFBn (blue) and iMGL (purple) peaks below. Peaks intersecting with GWAS risk variants (Risk peaks) are highlighted in red, and iMGL specific risk peaks are pink. Genes are at the bottom, with blue bars representing exons and black lines showing introns. The vertical line shows the location of rs6733839. Yellow highlight shows *BIN1*.

Modeling of ATAC peaks and risk colocalized eQTL effect in MGL included ten peaks at the *BIN1* locus. These candidate peaks contain variants associated with risk or were peaks whose magnitude is correlated with *BIN1* risk and are present in iMGL but not iFBn. There were no caQTL peaks that colocalized with risk at this locus. Visualization of this data together on a locus plot shows the GWAS signal for both AD and LBD overlap in a region upstream of the *BIN1* TSS (Figure 4C), these variants also have high eQTL effects in iMGL (purple), while the signal in ExN (blue) and InN (green) is low. We also found the peaks identified in our iFBn (blue bars) and iMGL (purple bars) each overlap significant GWAS SNPs, which we defined as risk peaks (red bars). Considering the *BIN1* colocalization signal is only found in iMGL we selected the risk peaks that only intersect with iMGL peaks for further functional study (Figure 4C, pink bars). This resulted in five peaks, two upstream, two intronic and one downstream that we hypothesized may be involved in transcriptional regulation of *BIN1* expression. Of these five that are present in iMGL but not iFBn, three were positively correlated with *BIN1* expression. Analysis of co-accessibility of peaks at the locus (Pliner *et al*., 2018) shows higher co-accessibility between peaks in iMGL relative to iFBn, including between the five peaks selected for functional analysis (Figure S8C).

### Silencing of BIN1 Risk-bearing Peaks Reveals Complex Transcriptional Control

Previous studies have demonstrated a 363-bp deletion containing AD-risk variant rs6733839 is sufficient to modulate microglial *BIN1* expression (Nott *et al*., 2019). However, it is unclear whether the effect of this deletion was limited to *BIN1* at this locus or whether the effect of deletion phenocopies transcriptional control induced by the GWAS lead SNP. Given that our iMGL analysis revealed five risk-bearing peaks around the *BIN1* locus, we aimed to increase the resolution at which these iMGL-specific putative enhancer regions could be functionally screened (Figure 5A).

**Figure 5.**
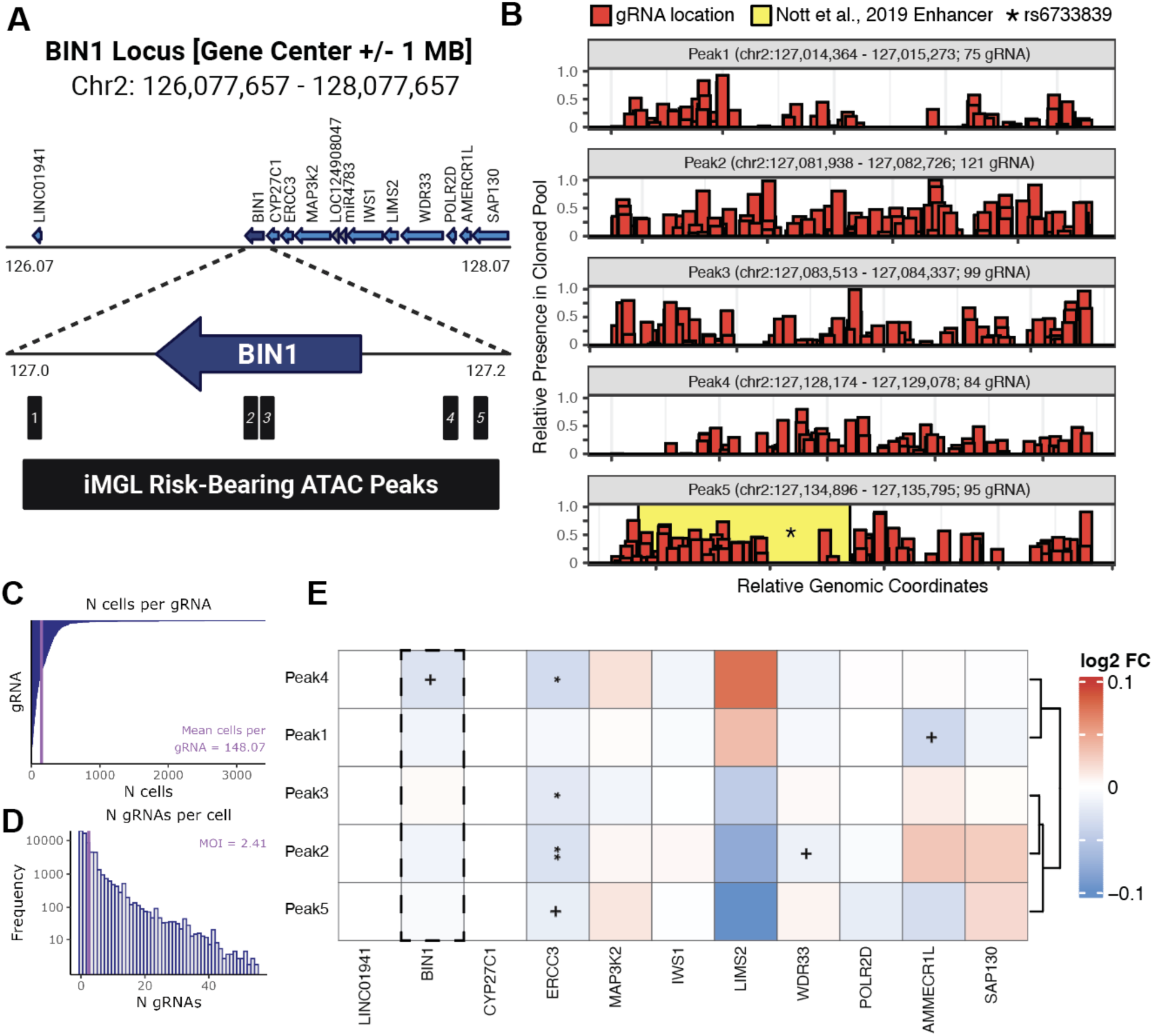
Silencing of BIN1 Risk-bearing Peaks Reveals Complex Transcriptional Control. A) Schematic of CRISPRi investigation of iMGL risk-bearing peaks at the *BIN1* locus. Risk peaks were within <100 kb of the *BIN1* TSS and hypothesized to control expression of genes within 1Mb of the *BIN1* gene center (13 genes met criteria, blue; LOC1290847 and miR4783 were not expressed in iMicroglia). B) All available gRNA within risk peaks were cloned into a lentiviral vector containing guide machinery. Median guide distribution of 474 targeting gRNAs in the final cloned plasmid pool reflects even targeting of peaks 1-5 (range = 75-121; average = 94.8 guides per peak). Lentiviral mediated expression in iMicroglia containing dCas9-KRAB (59,575 cells past quality control) resulted in C) an average of 148.07 cells per gRNA and D) an average MOI of 2.41. E) SCEPTRE analysis (“High MOI” version) of targeting and non-targeting (500 guides) gRNA revealed a dampening of *BIN1* with Peak-4 targeting guides (*p* < 0.1) and 4 peaks with a more drastic dampening of *ERCC3* expression. + *p* < 0.1, * *p* < 0.05, ** *p* < 0.01.

First, we designed a strategy to virally express guide RNAs in iMGL expressing dCas9-BFP-KRAB which would permit the epigenetic silencing of *BIN1* (Figure S9). In a pilot experiment, we introduced guides in the *BIN1* promoter, as well as guides we previously reported to control *LRRK2* expression in microglia (Langston *et al*., 2022), using lentiviral transduction of mature iMGL boosted by Vpx-Virus like particles (VLPs; Dolan et al., 2023) and showed that we could modify expression of target genes using this approach (Figure S10). Next, we performed the functional analysis screen in which we expressed a pool of 474 targeting guides distributed across the five risk-bearing peaks of interest (Figure 5B). Notably, there were no available Protospacer Adjacent Motifs (PAM) sequences resulting in 20bp guide binding sites overlapping with rs6733839, making it impossible to target the GWAS lead SNP directly. Therefore, we performed Single-Cell PerTurbation screens via conditional REsampling (SCEPTRE; Barry *et al*., 2024) with the “response_id” set to the gene expression of targets within 1 megabase of *BIN1* and “grna_target” set to the 5 risk-bearing peaks overlapping with AD and LBD risk at the *BIN1* locus (*See Methods*). This resulted in an experiment in which a total of 59,575 cells passed quality control at a mean cells per gRNA of 148.07 (Figure 5C) and an experimental MOI of 2.41 (Figure 5D). Cellwise-, pairwise-, and power-analysis of the 474 targeting and 500 non-targeting guide RNAs in our experiment indicated that perturbations could be detected without *p*-value inflation causing false discovery (Figure S11).

We identified a significant decrease (*p* < 0.05) in *ERCC3* when targeting 3 of the 5 peaks of interest (Figure 5E). While there is a caQTL for a peak and that peak is correlated with *ERCC3* and *BIN1* expression, the caQTL does not colocalize with AD genetic risk (Figure S12). Neither peak 5, which contains rs6733839 and the entirety of the previously reported 363-bp enhancer region (Nott et al., 2019), nor any other peak showed a statistically significant decrease in expression of *BIN1*; however, guides targeting peak 4 showed a non-significant trend towards decreased expression (*p* = 0.060). These findings are consistent with previous studies targeting regulatory elements upstream of *BIN1*, which also showed dampening of the expression of *ERCC3*, *IWS1*, and *MAP3K2* (Yang *et al*., 2023). Together, our CRISPRi screen at the *BIN1* locus suggest that microglia-specific risk-bearing enhancer elements bearing risk may have effects on nearby gene *ERCC3* in addition to previously reported effects on *BIN1*.

## Discussion

Here, we aimed to use QTL mapping to prioritize genes and variants associated with NDDs and complete functional validation *in vitro*. We focused on neurons and microglia, as these two cell types are thought to play major roles in NDD risk. Using a combination of data from iPSC-derived cells that we meta-analyzed with available brain data sets, we were able to identify many QTLs, several of which are relevant to NDDs, and performed a functional analysis of the *BIN1* locus that is relevant to multiple types of dementia.

We started with single cell expression and chromatin accessibility in a large number (*n*=138) of iPSC-derived microglia and forebrain neurons. Correlation of expression patterns with authentic brain cells was used to confirm the identity of the differentiated cell types and suggested that iPSC-derived cells are reasonable proxies for the transcriptional state of brain cells. Using this data to then generate QTL maps in iPSC-derived cells, we were able to identify numerically more events in iMGL than in iFBn.

We note that there is greater overlap in QTLs between iNeurons and developing brain compared to adult brain tissue. This result is consistent with our prior work (Bressan *et al*., 2023) that iPSC-derived neurons are closer to embryonic neurons than mature dopamine neurons. These results suggest that iFBn are relatively immature and attain a limited number of different transcriptional profiles compared to mature brain cells.

To overcome the relatively limited recovery of QTLs in iFBn, and to generally improve statistical power to detect such events, we then performed a meta-analysis between data from iPSC-derived cells and human brain data. With increased sample sizes for expression and genetic variability, we identified thousands of eQTL and caQTL across neurons and microglia. This QTL dataset was then intersected with GWAS from AD, PD and LBD to identify variants that may play a role in NDD pathogenesis by modifying gene expression or chromatin accessibility. Several key loci show evidence of association mediated by QTLs, including replication of our prior identification of QTL-driven expression at the *LRRK2* locus on Chr12 (Langston *et al*., 2022). The current analyses are therefore helpful in refining nominated QTLs to two major cell types. Together these data show that genetic risk is not simply a property of a variant or gene but the biological consequences depend on the cellular regulatory context in which it operates.

Using this prioritization method, we identified candidate regulatory peaks at one locus, *BIN1*, that is associated with both AD and LBD by GWAS, has MGL-specific eQTL and MGL-specific peaks that overlap risk loci. *BIN1* is promising candidate gene with predicted roles in inflammation (Sudwarts *et al*., 2022), autophagosome function (Palmer *et al*., 2025), and Tau spreading (Crotti *et al*., 2019), and several studies that have attempted to characterize the GWAS signal at the locus (Nott *et al*., 2019; Yang *et al*., 2023). We performed functional analysis in iMGL at the *BIN1* locus, using a Perturb-seq experiment with CRISPRi to block access to the putative *cis*-regulatory elements, and scRNA-seq to detect the expression of genes within the region. Similar approaches targeting non-coding regions containing genetic risk have previously been shown to elucidate microglia-specific risk for multiple sclerosis (Gallagher *et al*., 2025).

Our molecular dissection of the *BIN1* locus in this study revealed a more complex regulatory mechanism than previously thought. A prior study used CRISPR to delete an upstream enhancer element containing the AD and LBD risk variant rs6733839 and observed lower *BIN1* expression specifically in iPSC-derived microglia, but not neurons, astrocytes or stem cells, however they did not report expression of other nearby genes (Nott *et al*., 2019). In contrast, the Perturb-seq based method used in our study allowed for analysis of multiple regions across the locus and unbiased measurement of gene expression transcriptome wide. Using this approach, we nominated an alternate enhancer element which showed effects on expression of a second gene at this locus, *ERCC3*, which encodes the XBP protein involved in nucleotide excision repair (Weeda *et al*., 1991). These results were consistent with another study of predicted *cis*-regulatory elements in microglia, which showed that silencing using guides targeting an element upstream of *BIN1* also affected *ERCC3* and other genes at the locus (Yang *et al*., 2023). These published guides were located within our Peak4, where we observed a significant decrease in *ERCC3* expression as well as a suggestive effect on *BIN1* expression.

Our data also suggests that other risk-bearing regulatory elements at the *BIN1* locus (Peaks 2, 3, 5) similarly reduce *ERCC3* expression when targeted. While biallelic coding mutations in *ERCC3* are linked to Xeroderma pigmentosum B and Cockayne Syndrome (Cleaver *et al*., 1999; Hafsi and Saleh, 2026), for which the clinical presentation includes higher risk for dementia, very few studies have focused on its potential role in neurodegeneration. One study reported increased expression of ERCC3 protein in post-mortem brain tissue from Alzheimer’s disease patients compared to controls and hypothesized that this change is due to ongoing DNA damage during disease (Hermon *et al*., 1998), and more recent genetic evidence suggests that somatic mutations in genes including *ERCC3* may occur in the AD brain (Ivashko-Pachima *et al*., 2021). However, our analysis showed the GWAS signals in AD and LBD colocalized with *BIN1* and not *ERCC3*, suggesting that the reduction of *ERCC3* in this experiment is not associated with disease. Therefore, our data raise the possibility of complex *cis-* regulatory effects related to multiple genes at this locus and exhibit the importance of the endogenous genomic context in evaluation of disease risk.

### Limitations of the study

While iPSC-derived cells can be close transcriptional proxies of authentic brain cells, they remain relatively immature and with less diverse cell identities compared to rich variability in expression found in the adult brain. While this consideration is unlikely to be impactful for QTLs that are measurable throughout brain development, the younger epigenetic state of iPSC-derived cells will likely influence QTL patterns that are established after maturation or where aging acts as a modulatory factor.

Additionally, we note that cells in culture have distinct properties both from each other and from cells in the brain. Neurons in culture may attach relatively poorly to substrates and become fragile during dissociation whereas microglia are relatively robust. Such differences between how cell types behave in culture are likely to add technical noise to single cell genomics measures and thereby degrade detection of QTLs. Nuclear isolation from tissue may also be influenced by distinct technical variables that limit direct comparison between cell types. We also note that the cell types used here were chosen due to represent very different developmental and functional roles in the brain, with microglia developing outside of the CNS (Nayak, Roth and McGavern, 2014). The fact that we see very different sets of QTLs in these cell types supports this contention, but also means that many important cell types including astrocytes, oligodendrocytes, blood-brain barrier-associated cells and many specialized neuronal types, remain to be surveyed in future efforts.

## Data and Code Availability

Raw and processed anndata files for SCAT and SCRN are available on GEO under accession GSE335939. Summary results from *cis*-eQTL, *cis*-caQTL, gene-peak correlation, and colocalization with GWAS are available on Zenodo under project 15837842 (10.5281/zenodo.15837842). Parquet files from meta-analyzed microglia, inhibitory neurons and excitatory neurons are also available on Zenodo. Code for all processing and analyses are available on github (https://github.com/neurogenetics/ADRD_iPSC).

## Acknowledgements

This research was supported by the Intramural Research Program of the National Institutes of Health (NIH). The contributions of the NIH authors are considered Works of the United States Government. The contributions of the NIH author(s) are considered Works of the United States Government. The findings and conclusions presented in this paper are those of the authors and do not necessarily reflect the views of the NIH or the U.S. Department of Health and Human Services. This research was supported by the Intramural Research Program of the NIH, the National Institute on Aging (ZIAAG000931) and the National Institute of Neurological Disorders and Stroke (ZIANS003154). This work utilized the computational resources of the NIH HPC Biowulf cluster (http://hpc.nih.gov). Sequencing was completed by the NISC Comparative Sequencing Program, and the National Hearth Lung and Blood DNA Sequencing and Genomics Core.

## AMP PD Acknowledgement

Data used in the preparation of this article were obtained from the Accelerating Medicine Partnership® (AMP®) Parkinson’s Disease (AMP PD) Knowledge Platform. For up-to-date information on the study, visit https://www.amp-pd.org. The AMP® PD program is a public-private partnership managed by the Foundation for the National Institutes of Health and funded by the National Institute of Neurological Disorders and Stroke (NINDS) in partnership with the Aligning Science Across Parkinson’s (ASAP) initiative; Celgene Corporation, a subsidiary of Bristol-Myers Squibb Company; GlaxoSmithKline plc (GSK); The Michael J. Fox Foundation for Parkinson’s Research; AbbVie Inc.; Pfizer Inc.; Sanofi US Services Inc.; and Verily Life Sciences. ACCELERATING MEDICINES PARTNERSHIP and AMP are registered service marks of the U.S. Department of Health and Human Services.

## AMP PD Cohort Acknowledgements

Clinical data and biosamples used in preparation of this article were obtained from the (i) Michael J. Fox Foundation for Parkinson’s Research (MJFF) and National Institutes of Neurological Disorders and Stroke (NINDS) BioFIND study, (ii) Harvard Biomarkers Study (HBS) and the Stephen & Denise Adams Center for Parkinson’s Disease Research of Yale School of Medicine (CPDR-Y), (iii) National Institute on Aging (NIA) International Lewy Body Dementia Genetics Consortium Genome Sequencing in Lewy Body Dementia Case-control Cohort (LBD), (iv) MJFF LRRK2 Cohort Consortium (LCC), (v) NINDS Parkinson’s Disease Biomarkers Program (PDBP), (vi) MJFF Parkinson’s Progression Markers Initiative (PPMI), and (vii) NINDS Study of Isradipine as a Disease-modifying Agent in Subjects With Early Parkinson Disease, Phase 3 (STEADY-PD3) and (viii) the NINDS Study of Urate Elevation in Parkinson’s Disease, Phase 3 (SURE-PD3). BioFIND is sponsored by The Michael J. Fox Foundation for Parkinson’s Research (MJFF) with support from the National Institute for Neurological Disorders and Stroke (NINDS). The BioFIND Investigators have not participated in reviewing the data analysis or content of the manuscript. For up-to-date information on the study, visit michaeljfox.org/biofind. Genome sequence data for the Lewy body dementia case-control cohort were generated at the Intramural Research Program of the U.S. National Institutes of Health. The study was supported in part by the National Institute on Aging (program #: 1ZIAAG000935) and the National Institute of Neurological Disorders and Stroke (program #: 1ZIANS003154). The Harvard Biomarker Study (HBS) is a collaboration of HBS investigators [full list of HBS investigators found at https://www.bwhparkinsoncenter.org/biobank/] and funded through philanthropy and NIH and Non-NIH funding sources. The Stephen & Denise Adams Center for Parkinson’s Disease Research of Yale School of Medicine is funded through philanthropy and NIH and non-NIH funding sources. The HBS and CPDR-Y Investigators have not participated in reviewing the data analysis or content of the manuscript. Data used in preparation of this article were obtained from The Michael J. Fox Foundation sponsored LRRK2 Cohort Consortium (LCC). The LCC Investigators have not participated in reviewing the data analysis or content of the manuscript. For up-to-date information on the study, visit https://www.michaeljfox.org/biospecimens). PPMI is sponsored by The Michael J. Fox Foundation for Parkinson’s Research and supported by a consortium of scientific partners: [list the full names of all of the PPMI funding partners found at https://www.ppmi-info.org/about-ppmi/who-we-are/study-sponsors]. The PPMI investigators have not participated in reviewing the data analysis or content of the manuscript. For up-to-date information on the study, visit www.ppmi-info.org. The Parkinson’s Disease Biomarker Program (PDBP) consortium is supported by the National Institute of Neurological Disorders and Stroke (NINDS) at the National Institutes of Health. A full list of PDBP investigators can be found at https://pdbp.ninds.nih.gov/policy. The PDBP investigators have not participated in reviewing the data analysis or content of the manuscript. The Study of Isradipine as a Disease-modifying Agent in Subjects With Early Parkinson Disease, Phase 3 (STEADY-PD3) is funded by the National Institute of Neurological Disorders and Stroke (NINDS) at the National Institutes of Health with support from The Michael J. Fox Foundation and the Parkinson Study Group. For additional study information, visit https://clinicaltrials.gov/ct2/show/study/NCT02168842. The STEADY-PD3 investigators have not participated in reviewing the data analysis or content of the manuscript. The Study of Urate Elevation in Parkinson’s Disease, Phase 3 (SURE-PD3) is funded by the National Institute of Neurological Disorders and Stroke (NINDS) at the National Institutes of Health with support from The Michael J. Fox Foundation and the Parkinson Study Group. For additional study information, visit https://clinicaltrials.gov/ct2/show/NCT02642393. The SURE-PD3 investigators have not participated in reviewing the data analysis or content of the manuscript.

## Author Contributions

Conceptualization: MRC

Methodology: DJA, XR, AB

Software: N/A no new software

Validation: DJA

Formal analysis: SSA, JD, JRG, DJA

Investigation: XR, DJA, AB, SB, FH, GL, DMR, CAW, SS, MP

Resources: SWS, DJE, MK, LF, MRC

Data Curation: JD, JRG

Writing - original draft: XR, DJA, JRG, MRC

Writing - Review & editing: All

Visualization: XR, DJA, JRG

Supervision: SWS, MRC

Project Administration: DTW

Funding Acquisition: LF, SWS, MRC

## Methods

### Line selection

iPSC lines were generated from PBMCs banked as part of the Genetic and Epigenetic Signatures of Translational Aging Laboratory Testing Study (GESTALT; *n* = 89; IRB protocol 15-AG-0063) (Roy *et al*., 2021, 2023; Tsitsipatis *et al*., 2022, 2023; Reed *et al*., 2024), the Baltimore Study of Longitudinal Aging (BLSA; *n* = 8; IRB protocol # 03-AG-0325) (Ershler *et al*., 2005; Kuo *et al*., 2020, 2022; Olinger *et al*., 2025), and the NIH Movement Disorders Research Clinic (*n* = 41; IRB protocol: 01-N-0206)(Makarious *et al*., 2023). Consistent with the Declaration of Helsinki, written consent was obtained at the time of collection and all samples were deidentified prior to experimental work.

### iPSC growth and pooling

All iPSC lines were expanded in Essential 8™ medium (Thermo Scientific, A1517001) on Matrigel-coated (Corning, cat #354230) dishes, and Revita supplement (Thermo Scientific, cat # A2644501) was used for splitting and thawing. iPS cells were passaged using TrypLE™ Select Enzyme (Thermo Scientific, cat # 12605010). Growth rate of each iPSC line was observed and pools were determined based on similarity in growth rates. For each pool of six lines, an equal number of cells per iPSC line were combined. These pools were immediately used in iMGL or iFBn differentiation. iPSCs were frozen using Synth-a-Freeze™ Cryopreservation Medium (Thermo scientific, cat #A1254201).

### Microglia differentiation

Microglia were differentiated based on a previously published protocol (Brownjohn *et al*., 2018). On the first day of differentiation, six iPSC lines were mixed to create pools. We pooled 2,000 cells/iPSC line/well and plated a total of 12,000 cells/well in V-bottom shape 96-well ultra-low attachment plates (Sunoko) in Embryoid body medium (EBM) containing E8 media, 1x Revita, 50 ng/mL BMP-4, 20 ng/mL SCF, and 50 ng/mL VEGF-121. The next day, 80 μL of media/well was removed and replaced with 100 μL of the EBM without Revita. EBs were cultured for an additional 2 days, with EBM medium changed every day. On day 4, 24 EBs were transferred from 96-well plates to 1-well of the 6-well plate and cultured in 3 mL of Microglia progenitor media (MPM) containing X-VIVO 15 Lonza media, 2 mM GlutaMax, 55 mM beta-Mercaptoethanol, 100 ng/mL M-CSF, and 25 ng/mL IL-3. The EBs attached to the bottom of the plate within 3–4 days and began producing microglia progenitor cells between days 12 and 18. Floating microglia progenitor cells were collected by centrifugation (250 rcf, 5 min), and 0.6 million cells per well were seeded into non-coated 6-well plates for final differentiation in Microglia Maturation Media (MMM), which contains Advanced RPMI, 2 mM GlutaMax, 100 ng/mL IL-34, and 10 ng/mL GM-CSF. Cells were further differentiated for an additional 10 days, with full MMM media changes every other day, and were collected immediately after this period, at 30 days *in vitro*.

### Forebrain neuron differentiation

iPSCs were differentiated to forebrain neurons (FBn) as previously described with some modifications (Burkhardt *et al*., 2013; Reed *et al*., 2021). Briefly, iPSC lines were pooled as described above, plated in Matrigel-coated 6-well plates, and grown to confluence. FBn N3 differentiation media (50% DMEM/F12 (Thermo Fisher, 11320033), 50% Neurobasal (Thermo Fisher, 21103049) containing 0.5x GlutaMAX (Thermo Fisher, 25030-081), 1x Penicillin-Streptomycin (Thermo Fisher, 15140122), 0.5x B-27 minus vitamin A (Thermo Fisher, 12587010), 0.5x N2 supplement (Thermo Fisher, 17502048), 0.5x MEM Non-Essential Amino Acids (NEAA) (Thermo Fisher, 11140-050), 0.055 mM 2-mercaptoethanol (Thermo Fisher, 21985-023) and 1 µg/mL Insulin (Millipore Sigma, 91077C)) plus 1.5 µM Dorsomorphin (Tocris Bioscience, 3093) and 10 µM SB431542 (Stemgent, 04-0010-05) was added and refreshed daily until differentiation day 11. N3 without Dorsomorphin and SB431542 was replaced daily until day 16. From day 16 to 19, N3 media with 0.05 uM Retinoic acid (Sigma, R2625) was changed every day. Wells were split 1:2 on day 19 using Accutase (Sigma-Aldrich, A6964) and replated in N4 media (N3+RA) with ROCK inhibitor/ Y-27632 (Stemcell Technologies, 72302) on Poly-L-Ornithine (Sigma, 27378-049), Fibronectin (Fisher Scientific, CB40008A), Laminin (Sigma, L6274-.5MG), and Matrigel-coated plates. Media was changed to N4 the following day and every other day after until differentiation day 60. At day 60, cells were treated with Accutase for dissociation from the plate. Cells were gently pipetted and pelleted at 150 x *g*. Supernatant was aspirated and cells were washed in 0.04% BSA (Miltenyi Biotec, 130-091-376) in DPBS, and pelleted again at 150 x *g*. Supernatant was removed and cells were resuspended in 0.04% BSA and filtered through a 70 μm Flowmi cell strainer (Sigma, BAH136800070). Live cells were counted using a TC20 Automated Cell Counter (Bio-Rad, 1450102).

### Single-cell RNAseq and ATACseq library preparation and sequencing

Single cell gene expression and ATAC seq libraries were prepared independently from differentiated iFBn and iMGL cells as recommended by each protocol. Cells were dissociated, washed, filtered and resuspended at a final concentration of 1000 cells/uL and GEM generation targeting 10,000 cells was performed as directed for Chromium Next GEM 3’ Single Cell Kit v3.1 (10x Genomics, 1000120 and 1000121) or cells permeabilized and nuclei transposed following instructions from the Chromium Next GEM Single Cell ATAC Kit v1.1 (10x Genomics, 1000175 and 1000161).

Sequencing library preparation was performed as directed in the user manuals. Library concentration and sizes were determined using the Agilent 2100 Bioanalyzer with the High Sensitivity DNA kit (Agilent, 5067-4626). Paired-end sequencing was completed as recommended on a NovaSeq6000 with 28-10-10-91 paired-end reads for single-cell RNA-seq and 50-8-24-49 paired-end reads for single-cell ATAC-seq samples. Samples were processed using the count method from CellRanger (v5.0.1, introns included and refdata-gex-GRCh38-2020-A annotation file) and CellRanger-atac (v2.0.0, refdata-cellranger-arc-GRCh38-2020-A-2.0.0 annotation file) tool packages for RNA and ATAC, respectively. For ATAC data, the CellRanger-atac aggr command was used to combine all ATAC samples, including both cell-type differentiations so that all samples were quantified across a consistent peak feature set.

### Genotype preparation and imputation

Genotypes were called using GenomeStudio Genotyping Module (v.2.0, Illumina) with a GenCall threshold of 0.15. Sample and variant QC was done using the PLINK toolset (Purcell *et al*., 2007; Chang *et al*., 2015). Samples were excluded based on the following criteria: a genotype call rate < 95%, a mismatch between reported sex and genotypic sex, evidence of sample duplication (pi-hat > 0.8), or an excess of heterozygosity or inbreeding, defined as an F-statistic deviating by more than +/- 0.15. Prior to imputation, variants were excluded if they met any of the following criteria: monomorphic status, palindromic alleles, non-autosomal location, an overall missingness rate > 5.0%; a significant haplotype-based non-random missingness (*p* < 1.0×10^-4^); or deviation from Hardy-Weinberg equilibrium (*p* < 1.0×10^-10^). Genotype imputation was performed on the TopMed Imputation Server using the Trans-Omics for Precision Medicine (TopMed) reference panel (hg38). Phasing was conducted using Eagle v2.4, followed by imputation via minimac4 (Fuchsberger, Abecasis and Hinds, 2015; Das *et al*., 2016; Taliun *et al*., 2021). Post-imputation variants were retained only if they possessed an imputation quality score *R*^2^ > 0.8 and a minor allele frequency (MAF) > 0.01.

### Demultiplexing pooled samples

Demultiplexing of pooled samples was done with the demuxlet tool (Kang *et al*., 2018), the prepared subject genotypes (non-imputed) split by pool, converted to vcf format, and the aligned single-nuclei bam files were used to deconvolute the cells’ sample identities. The demultiplexing process was performed on the Google Cloud Platform (GCP) using the Cumulus/Demuxlet workflow (WDL, https://cumulus-doc.readthedocs.io/en/0.12.0/demuxlet.html), which executes the demuxlet tool contained in the Statgen Popcle suite (https://github.com/statgen/popscle). Job submission to GCP for execution was done via the Broad WDL runner (https://github.com/broadinstitute/wdl-runner) and GCP Life Sciences interface (https://cloud.google.com/life-sciences/docs). Scanpy (Wolf, Angerer and Theis, 2018) was used to read in the 10X filtered matrix files into an anndata object and integrate sample identity for the deconvoluted cells, along with sample information into the ‘obs’ information. For single-cell RNA data, this was done per sample and then combined all data into a single anndata object. Whereas for the single-cell ATAC data the integration of deconvoluted cell identity was done on the CellRanger-atac aggregated data.

### Clustering and cell-type identification

SCANPY was used to combine (RNA), filter, normalize, batch correct, and cluster the single-cell separately for each modality (RNA and ATAC) and each differentiated cell-type (FB and MG). Basic filtering was done with scanpy to exclude cells that did not show at least 200 genes in RNA or 1% of peaks in ATAC, genes that were not present in at least three cells, and cells that had more than 10% mitochondrial content (McCarthy *et al*., 2017). Counts were transformed to a total-count normalization of 10,000 reads per cell and log transformation. Cell type assignments were done using a modified version of the marker-based automatic cell-type annotation (MACA) tool (Xu *et al*., 2022). The modification made to MACA was to switch the clustering algorithm from Louvain to Leiden (Levine *et al*., 2015; Traag, Waltman and van Eck, 2019). MACA was run using multiple Leiden resolutions of (0.3, 0.4, 0.5, 0.6) and numbers of neighbors (5, 10, 50, 100). Gene marker sets for cell types present in the human central nervous system were used with MACA to assign cell types, where the cell-type marker set from the public database of single-cell experiments PanglaoDB, or previous publications of human brain cell types (McKenzie *et al*., 2018; Franzén, Gan and Björkegren, 2019; Bakken *et al*., 2021). MACA was also used to assign cell-type identifiers to the single-cell ATAC data, where the same gene marker sets were used but based on the intersection of these genes’ promoters with ATAC peaks present in either of the differentiated cell types.

The top highly variable features were used for clustering, accounting for pool batch and using a dispersion-based method by setting the flavor to ‘seurat’ (Satija *et al*., 2015); 6,000 genes for RNA and 18,000 peaks for ATAC. Each feature was scaled to unit variance with a max value of 10. Principal components analysis was used to reduce the dimensionality of the data from the top high variable features (genes or peaks) to 20 principal components (PC). Batch effects were corrected using the batch-balanced k-nearest neighbors (BBKNN) method (Polański *et al*., 2020) with the pools used as batches. The Leiden algorithm was used to identify clusters on the neighborhood graph; multiple resolutions were used (0.3, 0.45, 0.6). Differential expression for each cluster against all others, along with known marker genes for broad central nervous system cell types, was used to make the initial inferences for each cluster’s putative cell-type. The differential expression analysis of the clusters was performed with SCANPY using Wilcoxon rank-sum and Bonferroni multiple test correction. Uniform Manifold Approximation and Projection (UMAP) (McInnes *et al*., 2018) was used to visualize, inspect, and evaluate the clustering and initial MACA-based cell-type assignments.

Inspection and evaluation of the clusters and their cell-type assignments were performed using dendrograms, dot plots, and UMAP scatter plots over gene markers common to neurons and microglia, as well as sample attributes. After evaluation, a Leiden resolution of 0.3 was used for both cell-type differentiations and both modalities. Based on visual inspection of UMAP and dot plots, clusters of cells that appeared substantially different from the targeted differentiated cell types were excluded from further analysis.

### Pseudo Bulk conversion and trait preparation

After exclusion of non-target differentiated cell-type clusters, the per-cell data were converted to pseudo bulk using mean expression per donor from the normalized and log-transformed cell quantification. Conversion to pseudo bulk is a method that allows for overcoming biases of pseudoreplication and zero-inflation that impact analyses in single-cell experiments (Zimmerman, Espeland and Langefeld, 2021; Murphy and Skene, 2022). During conversion, the fraction of cells per Leiden cluster per donor was computed to retain as covariates, as well as the total cell count per donor per cell-type differentiation and modality. Additional covariates were computed based on means of per-cell values for each donor that were generated during the scanpy clustering, including: n_genes_by_counts, total_counts, total_counts_mt, pct_counts_mt, and n_genes. Post pseudo bulk conversion, any sample with a total cell count of 10 or less was excluded from further analysis. The cutoff of 10 cells was based on inspection of the per-sample cell count distributions by cell-type differentiation and modality. Based on this cutoff, 8 iFBn SCAT, 14 iFBn SCRN, 3 iMGL SCAT, and 3 iMGL SCRN samples were excluded.

Modality feature detection rates were computed, and any feature not detected in at least 25% of samples for a cell-type and modality was excluded from further analysis. Based on this detection threshold, 46,513 iFBn SCAT, 11,680 iFBn SCRN, 70,470 iMGL SCAT, and 17,838 iMGL SCRN features were included in analyses. All features were scaled using scikit-learn preprocessing MinMaxScaler. The data was evaluated based on global variance for known and unknown sources of technical variance and then corrected for these variance components. The top 25% of autosomal features by variance per modality were used to evaluate the global variance: 11,628 iFBn SCAT, 2,920 iFBn SCRN, 17,618 iMGL SCAT, and 4,460 iMGL SCRN.

Dimensionality reduction via Principal Component Analysis (PCA) was used to reduce these high variance features per modality into a smaller set of variance components. The number of components used for PCA was based on the maximum curvature of the compression accuracy based on R-squared (*R^2^*) and root mean squared error (RMSE) metrics using the Kneed package KneeLocator function. For the iFBn cell-type, 25 and 18 components were selected for SCAT and SCRN, respectively. While for the iMGL cell-type, 19 and 10 components were selected for SCAT and SCRN, respectively. To understand how PCA variance components may be related to known covariates, these components were modeled as targets of the known covariates. This variance component modeling was carried out using a generalized linear model (GLM from the statsmodels package). For iFBn SCAT the 25 PCA variance components had a reconstruction *R^2^* accuracy of 64.4% and a RMSE of 0.1198. Where the first PCA variance component accounted for 9.5% of the variance in the iFBn SCAT features used for modeling. This first PCA variance component was most strongly correlated with the known covariate ‘n_genes_by_count’. For iFBn SCRN the 18 PCA variance components had a reconstruction *R^2^* accuracy of 78.2% and a RMSE of 0.1043. Where the first PCA variance component accounted for 52.5% of the variance in the iFBn SCRN features used for modeling. This first PCA variance component was most strongly correlated with the known covariate ‘n_genes_by_count’. For iMGL SCAT the 19 PCA variance components had a reconstruction *R^2^* accuracy of 61.6% and a RMSE of 0.1157. Where the first PCA variance component accounted for 23.1% of the variance in the iMGL SCAT features used for modeling. This first PCA variance component was most strongly correlated with the known covariate ‘leiden_2’ cluster. For iMGL SCRN the 10 PCA variance components had a reconstruction *R^2^* accuracy of 84.1% and a RMSE of 0.0861. Where the first PCA variance component accounted for 52.5% of the variance in the iMGL SCRN features used for modeling. This first PCA variance component was most strongly correlated with the known covariate ‘leiden_4’ cluster.

In general the PCA variance components correlated with other known covariates that would be expected to be a source of variance such as pool number and Leiden cluster. During the modeling of each feature and variance components the model score was checked. For iFBn SCAT, the PCA variance components on average accounted for 67.8% of a feature’s variance. For iFBn SCRN, the PCA variance components on average accounted for 70.1% of a feature’s variance. For iMGL SCAT the PCA variance components on average accounted for 64.7% of a feature’s variance. For iMGL SCRN, the PCA variance components on average accounted for 65.5% of a feature’s variance.

Preparation of broad cell types from post-mortem ROSMAP DLPFC for donors without or with mild MCI followed a similar methodology based on the pseudobulk mean of the lop1p(CPM) cell data per donor. Like the differentiated cell types PCA was used to model and generate variance components per broad cell-type. Where the broad cell types prepared for *cis*-eQTL analysis included: astrocytes (Ast), endothelial (Endo), excitatory neurons (ExN), inhibitory neurons (InN), microglia (MGL), oligodendrocytes (Oli), and oligodendrocyte precursor cells (OPC). The number of variance components selected per post-mortem broad cell types ranged from 14 to 27. The reconstruction accuracy of these selected variance components had a *R^2^* range from 82.3% to 93.4%, and root mean squared error (RMSE) range from 0.1454 to 0.2993. Where the Endo cell-type was an outlier with an *R^2^* of 56.6% and RMSE of 0.6508.

### cis-QTL analyses

The pseudobulk converted features were formatted as phenotypes into a BED format to be used as traits for *cis* expression or chromatin accessibility quantitative trait locus (*cis*-eQTL and caQTL) analyses using tensortQTL (Ongen *et al*., 2016; Taylor-Weiner *et al*., 2019). Nominal *cis*-QTL analyses were run for autosomal features and their *cis* proximal variants within +/- 1Mb from the TSS of a gene and a minor allele frequency (MAF) of at least 5%. The *cis*-QTL regression analysis between variant genotype and the pseudobulk quantified features also included sex, the first six genetic population structure principal components, cell count, donor institute source, and the PCA variance components as covariates. Empirical *cis*–QTL analyses were carried out based on 10,000 permutations to compute empirical *p*-values. To adjust for the number of features tested, a Benjamini and Hochberg false discovery rate (FDR) method was applied to the empirical *p*-values. For iFBn SCAT autosomal *cis*-caQTL was based on 129 samples, 47,945 features (traits or ATAC peaks), and 10,168,161 variants, and no feature met statistical significance based on feature-wide FDR correction. For iFBn SCRN, autosomal *cis*-caQTL was based on 125 samples, 12,091 features (traits or genes), and 10,168,161 variants: 27 features were statistically significant based on feature-wide FDR correction. For iMGL SCAT, autosomal *cis*-caQTL was based on 133 samples, 68,984 features (traits or ATAC peaks), and 10,168,161 variants: 2,125 features were statistically significant based on feature-wide FDR correction. For iMGL SCRN, autosomal *cis*-caQTL was based on 133 samples, 17,252 features (traits or genes), and 10,168,161 variants: 2,211 features were statistically significant based on feature-wide FDR correction. For the cis-eQTL analysis of broad cell types from post-mortem ROSMAP DLPFC for donors without or with mild MCI, based on the clinical consensus diagnosis of cognitive status at time of death, the per cell-type analysis samples ranged from 206 to 211, features analyzed ranged from 10,082 to 21,086, and detected eQTL ranged from 568 to 4,568. Where the endothelial result set was an outlier based on 124 samples, 5,728 features analyzed, and 18 features with an eQTL detected.

### Correlation between genes and cis peaks

Correlations between genes and their *cis-*proximal ATAC peaks were also performed using tensorQTL. Like *cis*-QTL, the proximal distance was +/- 1Mb between the ATAC peaks and a gene transcription start site. In these regressions, the quantified values of each ATAC peak were the independent or exogenous variable, and the expression of the gene was the dependent or endogenous variable. The same set of covariates as used in the *cis*-eQTL analysis were used here: sex, the first six genetic population structure principal components, cell count, donor institute source, and the PCA variance components as covariates. Empirical correlation analyses were carried out based on 10,000 permutations to compute empirical *p*-values. To adjust for the number of genes tested, a Benjamini and Hochberg false discovery rate (FDR) method was applied to the empirical *p*-values. For iFBn SCAT and SCRN, *cis* correlations were based on 125 samples, 47,983 peaks, and 12,106 genes: 5,136 genes had one or more proximal peaks with a statistically significant correlation after FDR correction. For iMGL SCAT and SCRN, *cis* correlations were based on 134 samples, 69,024 peaks, and 17,283 genes: 11,096 genes had one or more proximal peaks with a statistically significant correlation after FDR correction.

### Meta-analyses of cis-eQTL studies

Meta-eQTL analysis was carried out between the eQTL results within this study and publicly available eQTL results from microglia. Three meta-analyses were performed based on the eQTL from excitatory neurons, inhibitory neurons, and microglia. For the meta-eQTL analysis of neurons, the eQTL studies include this study with differentiated neurons from 133 samples and excitatory (211 samples) or inhibitory (207 samples) neurons from post-mortem cells from 248 donors in the publicly available ROSMAP post-mortem Dorso-lateral Pre-frontal Cortex (DLPFC) datasets from three studies (Fujita *et al*., 2024). The meta-eQTL analysis of microglia included the results from this study based on differentiated microglia from 133 samples, microglia from 206 post-mortem ROSMAP DLPFC samples, and 391 post-mortem subjects from a previous meta-eQTL in microglia from Kosoy et al (Kosoy *et al*., 2022), which included microglia eQTL studies from two prior studies (Young *et al*., 2021; Lopes *et al*., 2022). A fixed effects meta-analysis using the eQTL summary statistics was carried out using METAL (Willer, Li and Abecasis, 2010), where study weight was based on sample size. A Benjamini and Hochberg false discovery rate (FDR) method was applied to the resulting *p*-values. While cell-type *cis*-eQTL summary statistics based on 192 post-mortem mixed region cortical samples from the Bryois et al study (Bryois et al., 2022) are also publicly available, these data were not included in the meta-eQTL analysis for this study. This study also includes samples from the ROSMAP DLPFC cohort with an unclear intersection of donors. Based on the selection criteria in this study and within the Bryois study, up to 20% of the samples may intersect. With this degree of possible overlap the inclusion of those summary eQTL statistics would have significantly biased the meta-eQTL analysis results within the current study. However, the Bryois et al results were still used to interrogate and inspect specific results within this study (see Figure 4B).

### Intersection and colocalization with common neurodegenerative disease risk

Prior to colocalization analysis, *cis*-QTL that may intersect with Alzheimer’s (Bellenguez *et al*., 2022), Lewy Body Dementia (Chia *et al*., 2021; Kaivola *et al*., 2023), or Parkinson’s disease (Nalls *et al*., 2019) risk were identified, based on whether a GWAS index risk variant displayed a nominal QTL *p*-value was 0.01 or less. Colocalization analysis was performed for these potential intersections of QTL and risk. The analysis methods used to determine colocalization between QTL results and disease risk was based on the single causal variant colocalization method originally described for the Coloc tool (Giambartolomei *et al*., 2014). This method was modified from Python implementations available in OpenTargets (Mountjoy *et al*., 2021) and tensorQTL (Taylor-Weiner *et al*., 2019). For any QTL and PD risk colocalization where the H4 posterior probability was at least 0.5, this colocalization was considered to be supported. The ‘H4’ posterior probability represents the probability that both traits are associated and share a single causal variant (Wallace, 2021). In some analyses, downstream of the colocalization, results with H4 posterior probabilities less than 0.5 were included as not supported but possible colocalization of signals.

### Intersection of Disease Risk Variants with Open Chromatin Regions

Genome-wide association study (GWAS) summary statistics for AD (Bellenguez *et al*., 2022), PD (Nalls *et al*., 2019), and LBD (Chia *et al*., 2021; Kaivola *et al*., 2023), were downloaded. Index risk variants, as specified in the respective GWAS publications, were used to represent the risk loci. Linkage disequilibrium estimates *D’* and *r^2^* were computed for each index variant with other variants chromosome-wide using PLINK (v1.9). Reference genotypes based on 10,418 samples from the Accelerating Medicine Partnership® (AMP®) Parkinson’s Disease (AMP PD) were used as the reference panel for LD estimation (Iwaki *et al*., 2021). Only variants with a minor allele frequency MAF > 0.001 were included. The full list of possible risk variants is based on two association significance thresholds: any variant reaching a genome-wide significance threshold (*p*-value < 5.0×10^-8^) or any variant in LD with a significant index variant that also has a suggestive association significance threshold of (*p*-value < 1.0×10^-5^). The genomic intervals for the ATAC peak features from both the iFBn and iMGL and the possible risk variants were both converted into BED to find their genomic intersections using pybedtools. This intersection identified specific ATAC peaks that include one or more of the possible disease risk variants. These possible risk-associated regulatory peaks were exported as a BED file for subsequent downstream analyses and visualizations.

### CRISPRi Guide Design and Cloning

Guide RNAs (gRNAs) targeted five risk-bearing peaks were designed using the CRISPOR online tool (v5.2) (Concordet and Haeussler, 2018). All gRNA with an MIT Specificity Score (as reported by CRISPOR) above 50 within the bounds of the peaks-of-interest were selected for pooled cloning. Non-targeted gRNAs were selected from a previously published list designed for pooled designs (Replogle *et al*., 2022). For positive-control pilots, promoter-targeting guide RNAs versus *BIN1* were selected from USCS Genome Browser for specificity, strand bias, and distance to TSS or selected from a previously published *LRRK2*-targeting study with a similar design (Langston *et al*., 2022). Non-targeting (*n*=500), BIN1 targeting (*n*=474) and pilot (BIN1 TSS: *n*=6, LRRK2-targeting: *n*=4) were ordered as pooled DNA oligos from TwistBiosciences (San Francisco, CA, USA). Cloning was performed in two steps as previously reported (Tian *et al*., 2019). pMK1334 was a gift from Martin Kampmann (Addgene plasmid # 127965; http://n2t.net/addgene:127965; RRID:Addgene_127965). A summary of all guide sequences is available in the supplement (Table S7).

First, a gBlock containing guide machinery and a “holdover guide” was cloned into a lentiviral vector containing a GFP reporter (VectorBuilder, VB900088-2229upx; pLV[Exp]-CMV>EGFP) via KAPA HiFi HotStart ReadyMix (Roche, 07958927001). Following digestion of the plasmid with restriction enzymes SpeI (NEB, R3133 in rCutSmart^TM^ (NEB, B6004S) for 2 hours at 37°C, the digested backbone was gel-purified (Qiagen, 28706; QIAquick Gel Extraction Kit). The assembly was completed in 20 uL aliquots at a 3:1 (insert:vector) ratio for 1 hr at 50°C. The reactions were then pooled and purified via ethanol precipitation by pelleting the product with 6.6 µL of 3M Sodium acetate (ph 5.2), 0.3 µL of GlycoBlue and 180 µL of 100% ethanol. The product was washed three times by resuspending the pellet in ice cold 70% ethanol and centrifugating at 16,000 g for 10 minutes at 4°C. These products were stored at−20°C until ready for transformation. Transformation of pooled libraries was performed via electroporation of MegaX DH10B T1R Electrocomp^TM^ cells (ThermoFisher, C640003). Electroporation was performed with a BioRad MicroPulser Electroporator (BioRad #1652100) in 0.1 µm cuvettes with the manual setting ‘2.00’. Cells were recovered per manufacturer instructions, shaking at 225 RPM for 1 hour at 37°C before streaking on plates of LB 1.5% Agar (KD Medical; BLF-7070) containing ampicillin (100 µg/mL; ThermoFisher J60977.14). Colonies were picked and DNA was extracted via QIAfilter Plasmid Midi Kit (Qiagen, 12243) per manufacturer instructions. Colonies were sequenced via Oxford Nanopore long read sequencing (Quintara Bio, Cambridge, MA, USA; “Whole Plasmid Sequencing” service) to confirm successful insertion of guide machinery and compared to the theoretical sequence of the donor plasmid with machinery insertion. Colonies determined to have 100% alignment to the theoretical sequence were used for pooled guide cloning.

Second, pooled vectors were created via dual digestion around the “holdover guide” followed by pooled assembly with KAPA HiFi HotStart ReadyMix (Roche, 07958927001). Digestion of the plasmid containing guide machinery was performed with BstXI (NEB, R0113) and BlpI (R0585) in NEBuffer r.2.1 (NEB, B6002) as described above. Pooled assembly and transformation were performed as described above, with one modification to post-transformation cell amplification. Cells containing pooled assemblies were amplified overnight in Super Broth with Modified Phosphates (KD Medical, BLE-3160) containing ampicillin (100 µg/mL; ThermoFisher J60977.14). To assess presence of the “holdover guide” and distribution of sequences in each respective pool, guide distributions were confirmed via short read sequencing of the the guide machinery region amplified via polymerase chain reaction (Quintara Bio, Cambridge, MA, USA; “AmpExpress Amplicon Sequencing” service; Table S6). In short, Tguide distribution was performed by aligning reads (zero tolerance) to a custom reference of the amplicon region containing every possible guide insertion. Second step cloning reactions containing > 90% intended guides with minimal run-away (no guides containing >5X theoretical even distribution in the pool) were stored at−20°C indefinitely for viral production.

### Production of virus and virus-like particles for CRISPRi screen

Virus-like particles (VLP) containing Vpx (Kim *et al*., 2019) and lentivirus containing gRNA libraries from pooled cloning were produced in-house. Lenti-X™ 293T cells were plated and transfected at approximately 80% confluency using jetPRIME® per manufacturer protocol (1:2 DNA:reagent in 100 µL buffer). For the production of Vpx-VLPs, cells were co-transfected with 5 µg of a VSV-G envelope expressing plasmid (pMD2.G) and 5 µg of a Vpx expressing plasmid. pSIV-D3psi/delta env/delta Vif/delta Vpr was a gift from Jeremy Luban (Addgene plasmid #132928; http://n2t.net/addgene:132928; RRID:Addgene_132928). Forty-eight hours post-transfection, the cell culture supernatant containing the VLPs was collected, centrifuged, and filtered through a 0.2 µm filter. The unconcentrated supernatant was then aliquoted and flash-frozen for storage at−80°C. For the production of lentivirus containing gRNA pooled libraries, the same protocol was followed replacing the 5 µg envelope-to-5 µg donor plasmid ratio with a 5 µg:5 µg:5 µg ratio of VSV-G envelope (pDM2.G, addgene #12259), packaging plasmid (psPax2 addgene #12260), and gRNA pools. pDM2.G and psPAX2 were gifts from Didier Trono (Addgene plasmids #12259 and 12260; http://n2t.net/addgene:12259; RRID:Addgene_12259 and http://n2t.net/addgene:12260; RRID:Addgene_12260). In lieu of calculating viral titers, quality control of each viral batch was determined by a pilot in iMicroglia assessing GFP expression following transduction with a range of dilutions in M3 media (1:6, 1:12, 1:60, 1:120, 1:600, 1:6,000). To ensure even transduction across batches of virus, effective dose was determined by the lowest dilution of virus resulting in >80% GFP expression after 48 hours of transduction.

### Lentiviral transduction of iPSC-derived microglia

iPSC-derived microglia expressing dCas9-BFP-KRAB were transduced with the pooled gRNA lentivirus, boosted by co-administration of Vpx-VLPs. iMicroglia were produced as described above (see “Microglia differentiation”) from an iPSC line (genetic background: WTC11) containing dCas9-BFP-KRAB (Tian *et al*., 2019). pC13N-dCas9-BFP-KRAB was a gift from Martin Kampmann (Addgene plasmid # 127968; http://n2t.net/addgene:127968; RRID:Addgene_127968). Within 10 days of maturation, expression of the pooled library was performed as follows. Mature microglia were lifted off of uncoated plastic T75 flasks with Accutase (Sigma-Aldrich, A6964) and replated in 6-well dishes (1E6 cells per well) with fresh media containing unconcentrated Vpx-VLPs at a ratio of 10 µL per 10,000 cells. Twelve hours following replating in VLP-containing media, cells were transduced with unconcentrated virus at a ratio experimentally determined per batch. This resulted in a high multiplicity of infection (MOI) to ensure a high percentage of cells received a gRNA, as confirmed in a pilot experiment (see Supp. Figure 10) and experimentally determined via Single Cell 5’ Gene Expression and CRISPR Screening (GEX+CRISPR) sequencing.

### GEX+CRISPR experimental workflow

Following lentiviral transduction, a single-cell perturbation screen was conducted using the 10x Genomics Next GEM Single Cell 5’ Gene Expression and CRISPR Screening (GEX+CRISPR) platform. The transduced microglia were processed according to the manufacturer’s protocol to generate single-cell libraries for both gene expression and gRNA identity. In the initial discovery experiment, a total of 59,575 cells were sequenced and passed quality control filters. A subsequent validation experiment was performed with freshly prepared virus, yielding 43,648 cells that passed quality control.

iMGL for CRISPR studies were dissociated to single cells. The resulting single cell suspension was then prepared using the 10x Genomics Chromium Next GEM Single Cell 5’ HT kit v2 (10x Genomics, 1000374) and loaded on Single Cell Chip N (10x Genomics, 2000375). Single cell 5’ gene expression and CRISPR screening libraries were constructed as directed in the user guide (10x Genomics, CG000512 Rev. B). All libraries were quantified with Qubit 1X dsDNA HS reagents (Invitrogen, Q33231). The average fragment size was determined using high sensitivity DNA D5000 screentape analysis (Agilent Technologies, 5067-5593, 5067-5592). The gene expression library and CRISPR screening library were pooled at concentrations of 4 nM and 1 nM, respectively. Paired end sequencing was performed on one 10B flowcell (replicate 1; Illumina, 20085596) or two 1.5B flowcells (replicate 2; Illumina, 20104703) on a NovaSeqX with a read length of 26-10-10-90. BCL files were converted to fastq using BCL convert (Illumina, v4.2.7) on the NIH HPC.

### SCEPTRE Analysis of Locus-wide CRISPRi Effects

Raw fastq were first processed using 10x Genomics Cell Ranger software (10xGenomics, v8.0.0) where RNA features were aligned to hg38 (CellRanger reference 2024-A, GRCh38v110) with the addition of the GFP reporter sequence (VectorBuilder, VB900088-2229upx; pLV[Exp]-CMV>EGFP). Guide features were constructed from the CRISPOR output and used as “feature-ref” in cellranger’s count function. Technical replicates from the 10xGenomics library preparation were calculated separately and merged prior to proceeding with downstream analysis.

The resulting filtered feature-barcode matrices were imported into R and stored as a Seurat Object (Hao *et al*., 2024). The effects of guides targeting BIN1 risk peaks was tested against the expression of all genes within 1Mb of the center of the *BIN1* transcript (11 of 13 genes containing non-zero values in >10 cells). The analysis was performed using the SCEPTRE (Single-Cell Perturbation screens via conditional REsampling) R package (Barry *et al*., 2021, 2024). In brief, positive control pairs, consisting of gRNAs targeting the *BIN1* and *LRRK2* promoters and their respective genes, were explicitly defined. The gRNA assignment in individual cells was determined using the "mixture model" method within SCEPTRE. A calibration check was performed to confirm that the statistical model did not suffer from *p*-value inflation, which could lead to an elevated false discovery rate per author’s recommendation. Finally, the discovery analysis was run to identify significant regulatory relationships between the targeted peaks and nearby gene expression per default parameters in the “High MOI” SCEPTRE vignette.

## Supplemental Information

Document S1: Supplemental Figures S1-S12.

**Figure S1:** Cohort Demographics. A) Ancestry of donors for iPSC lines. B) Count of donors by age at PBMC collection split by sex (Male=black bars, Female=white bars).

**Figure S2:** QTL and AD risk colocalization at the *PLEKHA1* locus. A) *PLEKHA1* forest plot of eQTL results from each study and the overall combined effect for the meta-eQTL analyses. Bryois MGL is not included as part of the meta-analysis but shown for comparison. Plot shows regression statistics per study for the AD GWAS risk index variant. B) *PLEKHA1* locus plot showing AD risk colocalization for rs7908662 (vertical line) with eQTL signal and intersection with ATAC peaks in iMGL (purple bars) and iFBn (blue bars). The *PLEKHA1* coding region is highlighted in yellow. The pink bar shows a iMGL specific risk peak in an intron of *PLEKHA1*.

**Figure S3:** QTL and AD risk colocalization identifies risk peaks in neurons and iMGL. A) Locus plot around *PRSS36* with AD GWAS on top, QTL signal in the middle and peaks on the bottom. Blue bars show iFBn peaks, purple bars show iMGL peaks, red bars show peaks overlapping GWAS risk variants, and pink bars show iMGL specific peaks that overlap risk. B) Plot of the *ACE* locus. Pink bars show iMGL specific peaks overlapping AD risk.

**Figure S4:** QTL and PD risk colocalization at the *LRRK2* locus. A) *LRRK2* forest plot of eQTL results from each study and the overall combined effect for the meta-eQTL analyses. Plot shows regression statistics per study for the PD GWAS risk index variant. B) *LRRK2* locus plot showing PD risk colocalization for rs76904798 (vertical line) with eQTL and intersection with ATAC peaks in iMGL (purple bars) and iFBn (blue bars). The locus region is truncated on the right to avoid including the GWAS p-value for *LRRK2* G2019S, the inclusion of that high risk coding mutations squashed the GWAS *p*-values for the common complex risk locus. The *LRRK2* coding region is highlighted in yellow. The pink bars show five iMGL specific peaks overlapping PD risk variants.

**Figure S5:** QTL and PD risk colocalization at the *RAB29* locus. A) RAB29 forest plot of eQTL results from each study and the overall combined effect for the meta-eQTL analyses. Bryois MGL is not included as part of the meta-analysis but shown for comparison. Plot shows regression statistics per study for the PD GWAS risk index variant. Plot shows regression statistics per study for the PD GWAS risk index variant. A) Locus plot of *RAB29* (yellow box) and rs823118 (vertical line) showing PD GWAS risk, eQTL signal, and chromatin peaks identified by cell type.

**Figure S6:** QTL and PD risk colocalization at the *SPNS1* locus. A) *SPNS1* forest plot of eQTL results from each study and the overall combined effect for the meta-eQTL analyses. Bryois MGL is not included as part of the meta-analysis but shown for comparison. Plot shows regression statistics per study for the PD GWAS risk index variant. Plot shows regression statistics per study for the PD GWAS risk index variant for *SPNS1*. B) *SPNS1* (gray box) locus plot showing PD risk for rs2904880 (vertical line) and intersection with eQTL and iMGL peaks. The pink bar shows a iMGL specific peak overlapping PD risk variants at the 3’ end of *SPNS1*.

**Figure S7:** LBD risk genes from GWAS colored by colocalization in at least one microglia population with eQTL H4 score (darker red is more significant). Results are filtered for protein coding genes only, full results are in Table S3.

**Figure S8:** BIN1 expression and locus characterization. A) Boxen/Scatter plot of BIN1 gene expression level by genotype in iMGL per allele dosage for the AD/LBD risk index variant rs6733839. B) BIN1 expression by genotype in single-nucleus microglia data from the DLPFC from the ROSMAP cohort. C) *BIN1* locus tracks figure for co-accessibility of open chromatin peaks relative to cell-type and peaks containing associated risk variants. Blue arcs represent co-accessibility in iFBn and purple arcs show co-accessibility in iMGL. Shaded regions highlight co-accessibility in regions where peaks were selected for functional analysis.

**Figure S9:** Cloning strategy and QC. A) Guide machinery inserted into lentiviral plasmid with GFP reporter (pLV[Exp]-CMV>EGFP; VectorBuilder VB9000088-2229upx) in two steps. First, guide machinery with a holdover guide (scramble) inserted into the lentiviral vector (See Methods, Table S6). Second, pooled cloning removes the holdover guide and inserts guides of interest (Table S7). B) Distribution of pooled gRNAs for BIN1 peak-targeting experiment measured by amplicon sequencing of plasmid pool following cloning step 2 (See Methods, Table S6). Intended guides determined by gRNA sequences ordered for cloning (Table S7). Present guides determined by at least 1 read from amplicon sequencing aligning with 0% mismatch to intended guide sequence. Acceptable guides determined by at least 50% intended relative abundance in the pool (i.e. 1 guide in 10 guide pool = 10% theoretical abundance, “acceptable” if measured abundance > 5%). C) Measured relative abundance of peak-targeting guides in amplicon sequencing (dotted: theoretical abundance, solid: “acceptable” cutoff for relative abundance).

**Figure S10:** LRRK2 and BIN1 promoter pilot. Target-feature correlation performed in SCEPTRE with “High-MOI” default parameters (See Methods). Cell-wise A) and pairwise B) quality control metrics reveal appropriate cutoffs for guides-in-cell and guides-per-target. C-F) Representative distribution of guide counts per cell. G-I) Target-feature association quality control metrics as provided by SCEPTRE. J) Fold change in feature versus collective action of targeted guides (*** *p*_unadjusted < 0.001, ** *p*_unadjusted < 0.01).

**Figure S11:** SCEPTRE Analysis of candidate enhancer inhibition. Target-feature correlation performed in SCEPTRE with “High-MOI” default parameters (See Methods). Cell-wise (A) and pairwise (B) quality control metrics reveal appropriate cutoffs for guides-in-cell and guides-per-target. C) Representative distribution of targeting versus non-targeting guide counts per cell. D-G) Target-feature association quality control metrics as provided by SCEPTRE. H) Fold change in feature versus collective action of targeted guides (** *p*_unadjusted < 0.01,* *p*_unadjusted < 0.05, + *p*_unadjusted < 0.1).

**Figure S12:** caQTL correlated with ERCC3 and BIN1 expression does not colocalize with genetic risk of AD. Forest plot in different MGL populations shows no colocalization of ERCC3 with AD risk.

## Supplemental Tables

**Table S1:** Donor line demographics and pooling strategy for iMicroglia and iNeuron differentiations.

**Table S2:** Single cell demultiplexing by genotype summary.

**Table S3:** Colocalization scores for eQTL and risk in AD, PD, and LBD.

**Table S4:** Colocalization scores for caQTL and risk in AD, PD, and LBD.

**Table S5:** Peak-gene association and risk scores for AD, PD, and LBD.

**Table S6:** Plasmids and oligos used for CRISPRi study.

**Table S7:** gRNA sequences used for CRISPRi study.

