## Supplemental Figures for "Integration of cell-specific gene expression and chromatin accessibility facilitates localization of neurodegenerative risk in microglia"

### Supplementary Figures

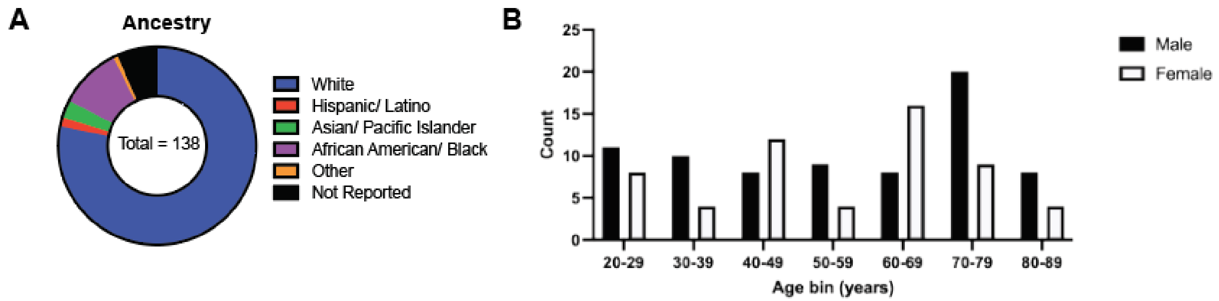


**Supplementary Figure 1.** Cohort Demographics. A) Ancestry of donors for iPSC lines. B) Count of donors by age at PBMC collection split by sex (Male=black bars, Female=white bars).

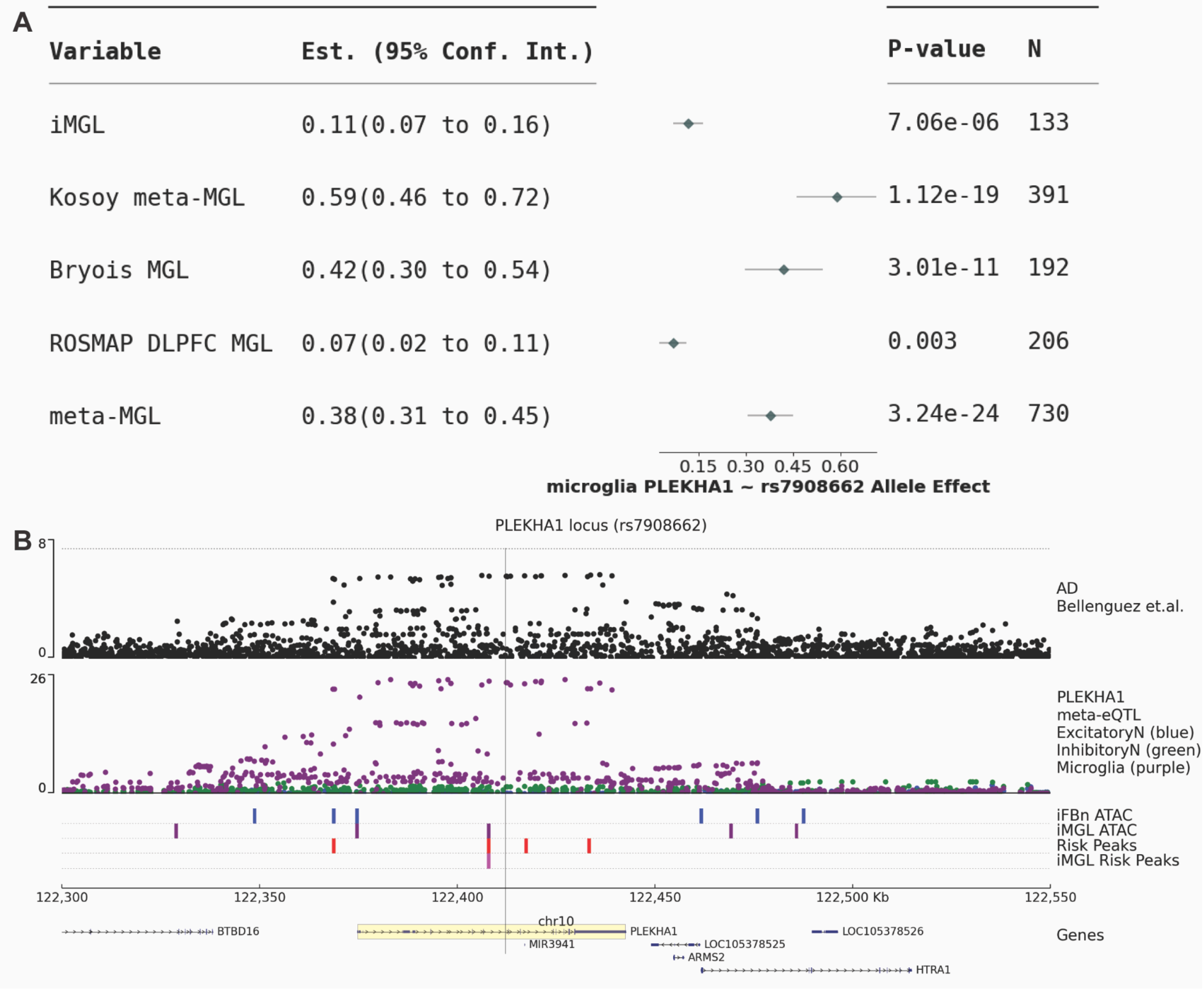


**Supplementary Figure 2:** QTL and AD risk colocalization at the *PLEKHA1* locus. A) *PLEKHA1* forest plot of eQTL results from each study and the overall combined effect for the meta-eQTL analyses. Bryois MGL is not included as part of the meta-analysis but shown for comparison. Plot shows regression statistics per study for the AD GWAS risk index variant. B) *PLEKHA1* locus plot showing AD risk colocalization for rs7908662 (vertical line) with eQTL signal and intersection with ATAC peaks in iMGL (purple bars) and iFBn (blue bars). The *PLEKHA1* coding region is highlighted in yellow. The pink bar shows a iMGL specific risk peak in an intron of *PLEKHA1*.

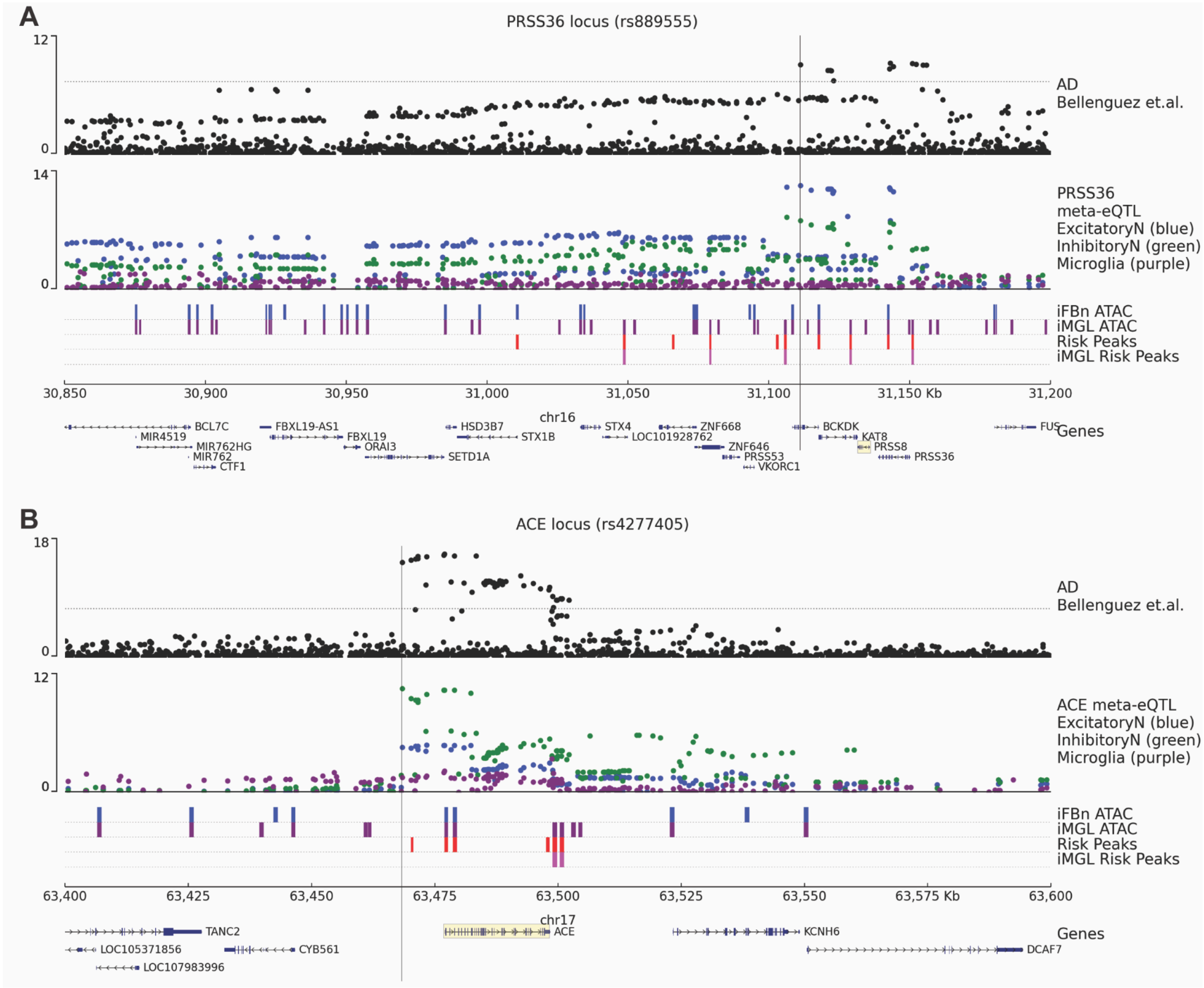


**Supplementary Figure 3:** QTL and AD risk colocalization identifies risk peaks in neurons and iMGL. A) Locus plot around PRSS36 with AD GWAS on top, QTL signal in the middle and peaks on the bottom. Blue bars show iFBn peaks, purple bars show iMGL peaks, red bars show peaks overlapping GWAS risk variants, and pink bars show iMGL specific peaks that overlap risk. B) Plot of the ACE locus. Pink bars show iMGL specific peaks overlapping AD risk.

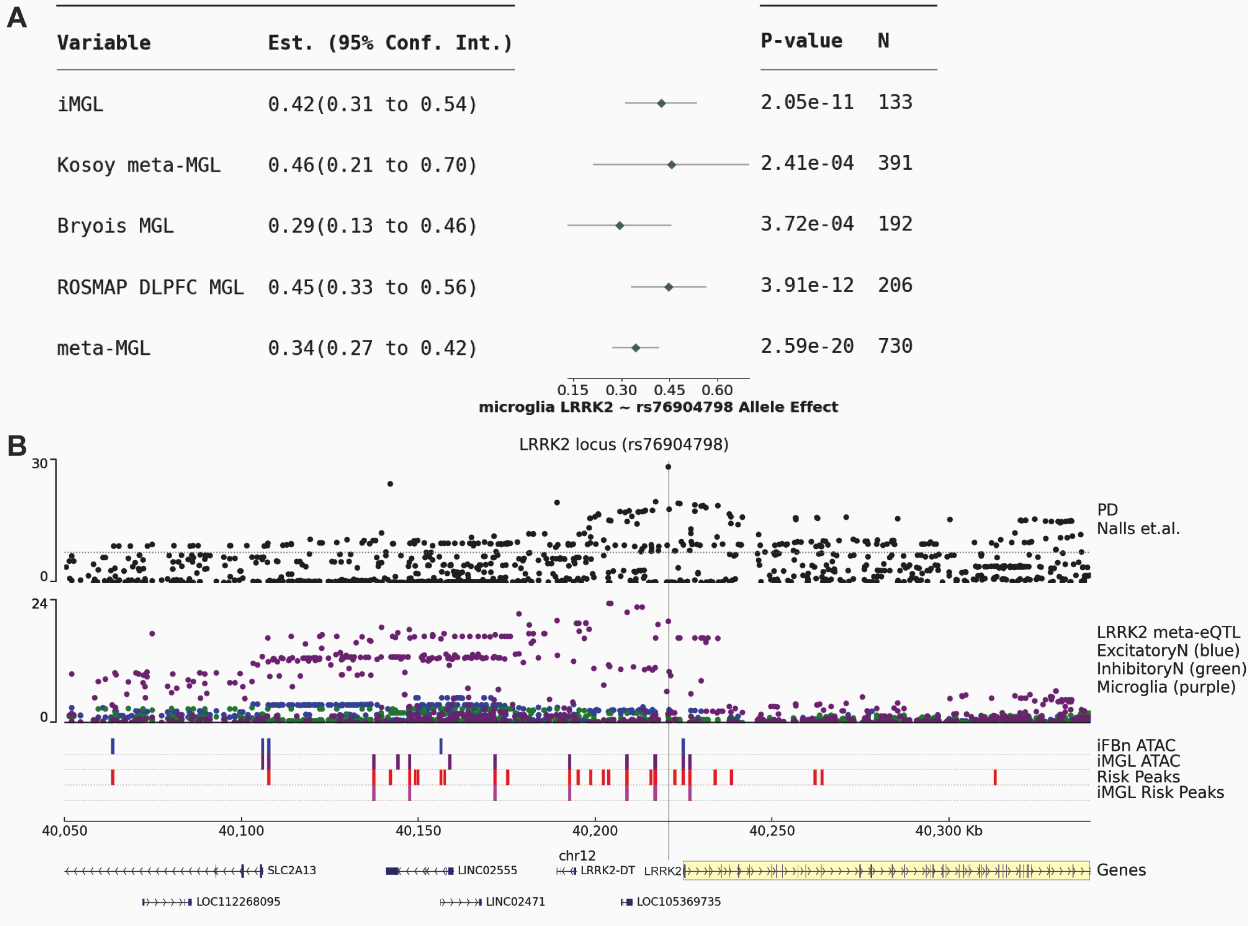


**Supplementary Figure 4:** QTL and PD risk colocalization at the *LRRK2* locus. A) *LRRK2* forest plot of eQTL results from each study and the overall combined effect for the meta-eQTL analyses. Plot shows regression statistics per study for the PD GWAS risk index variant. B) *LRRK2* locus plot showing PD risk colocalization for rs76904798 (vertical line) with eQTL and intersection with ATAC peaks in iMGL (purple bars) and iFBn (blue bars). The locus region is truncated on the right to avoid including the GWAS p-value for *LRRK2* G2019S, the inclusion of that high risk coding mutations squashed the GWAS p-values for the common complex risk locus. The *LRRK2* coding region is highlighted in yellow. The pink bars show five iMGL specific peaks overlapping PD risk variants.

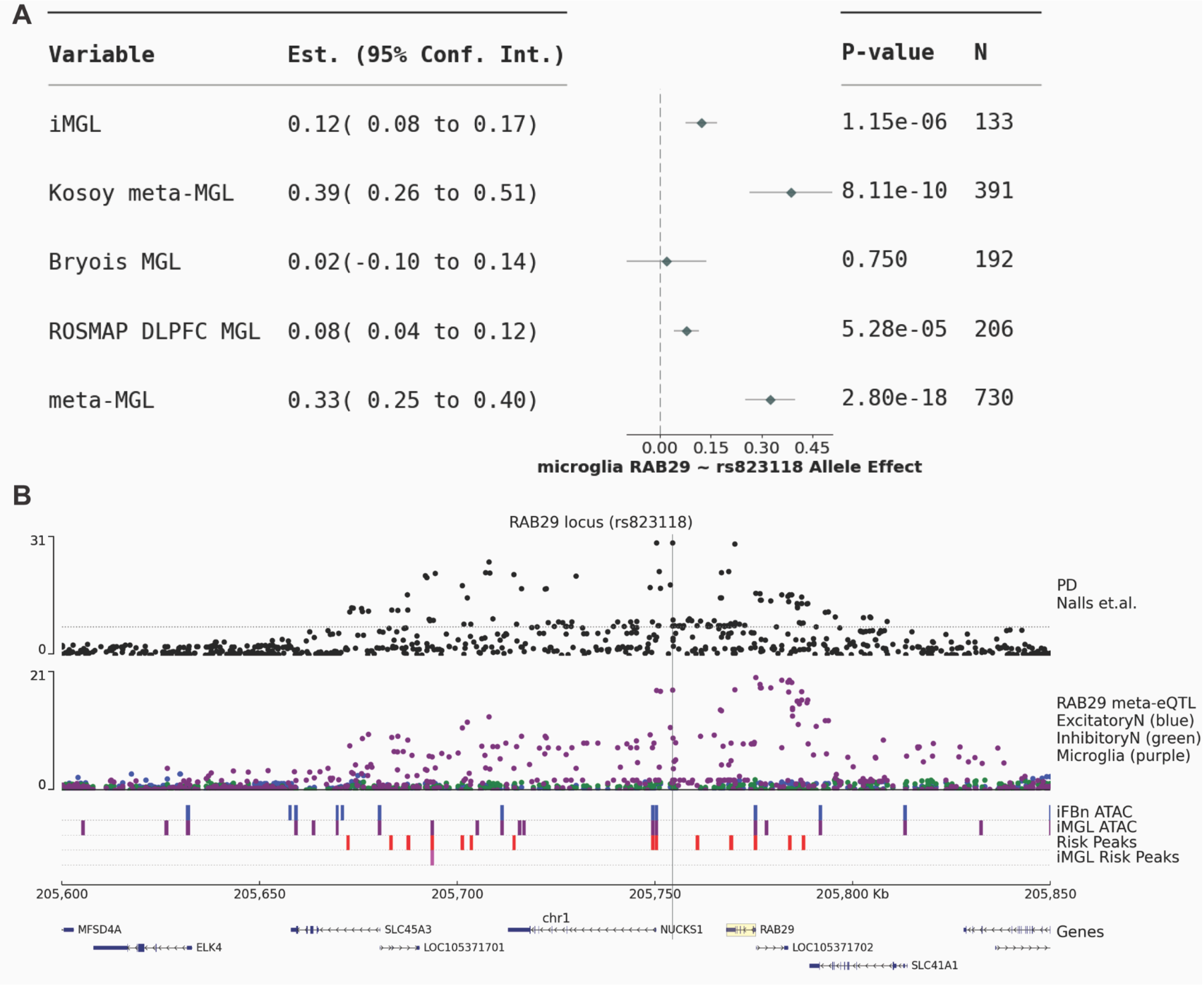


**Supplementary Figure 5:** QTL and PD risk colocalization at the *RAB29* locus. A) RAB29 forest plot of eQTL results from each study and the overall combined effect for the meta-eQTL analyses. Bryois MGL is not included as part of the meta-analysis but shown for comparison. Plot shows regression statistics per study for the PD GWAS risk index variant. Plot shows regression statistics per study for the PD GWAS risk index variant. A) Locus plot of *RAB29* (yellow box) and rs823118 (vertical line) showing PD GWAS risk, eQTL signal, and chromatin peaks identified by cell type.

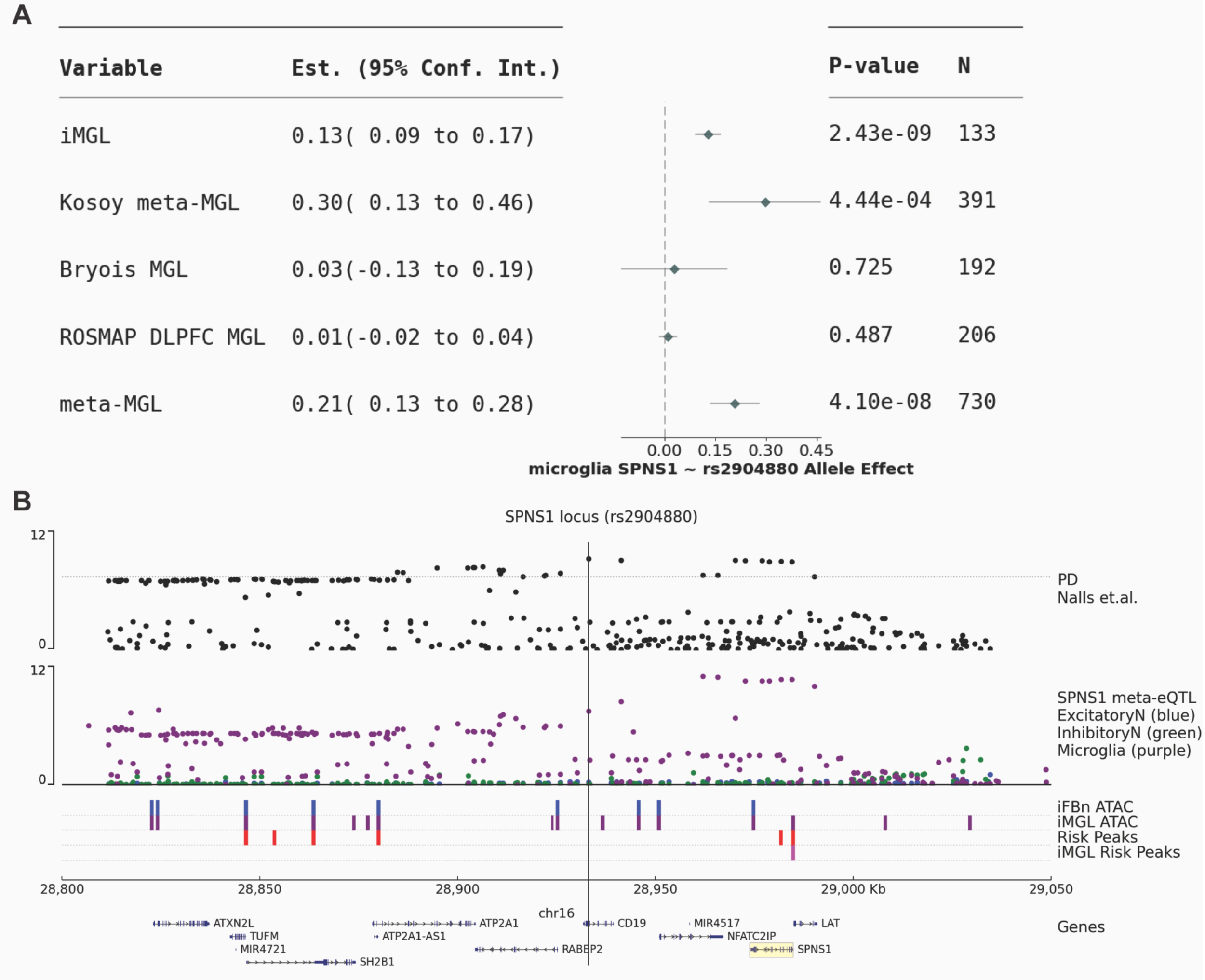


**Supplementary Figure 6:** QTL and PD risk colocalization at the *SPNS1* locus. A) *SPNS1* forest plot of eQTL results from each study and the overall combined effect for the meta-eQTL analyses. Bryois MGL is not included as part of the meta-analysis but shown for comparison. Plot shows regression statistics per study for the PD GWAS risk index variant. Plot shows regression statistics per study for the PD GWAS risk index variant for *SPNS1*. B) *SPNS1* (gray box) locus plot showing PD risk for rs2904880 (vertical line) and intersection with eQTL and iMGL peaks. The pink bar shows a iMGL specific peak overlapping PD risk variants at the 3' end of *SPNS1*.

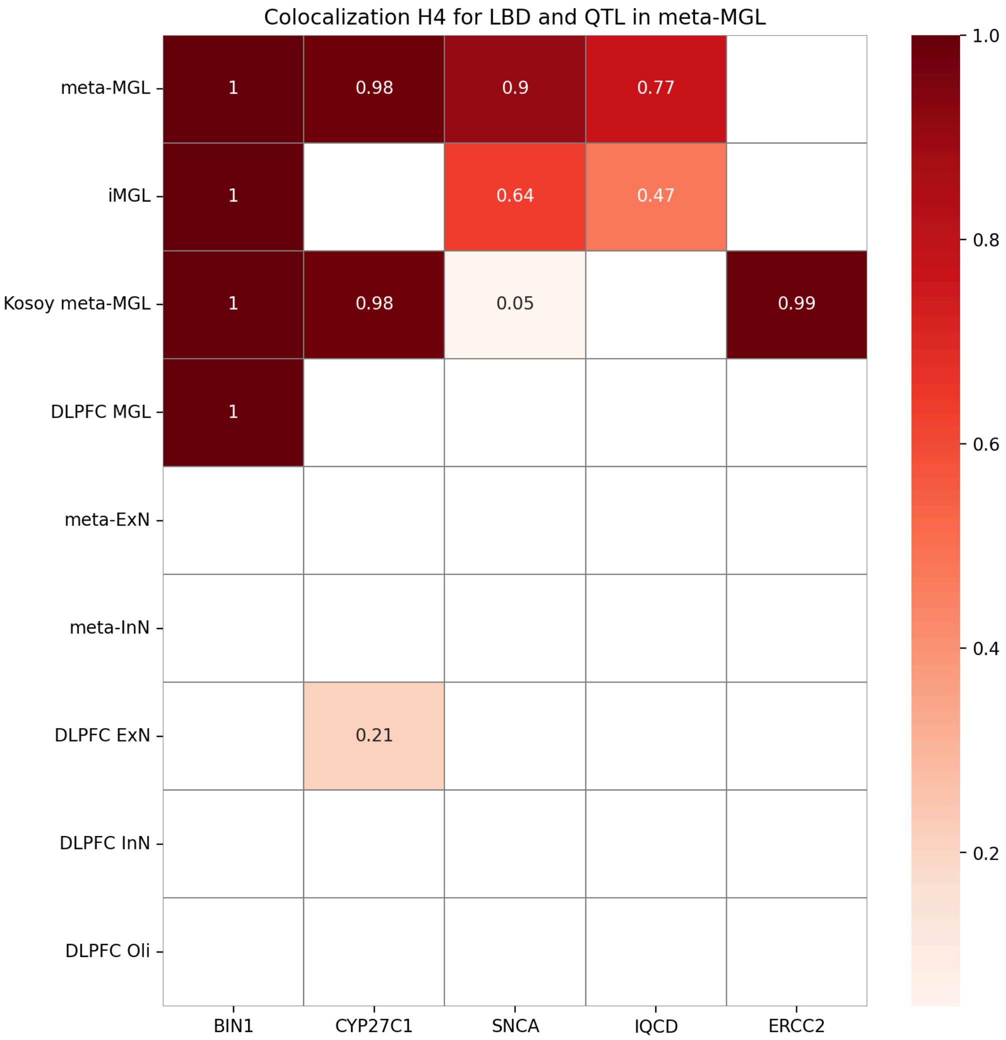


**Supplementary Figure 7:** LBD risk genes from GWAS colored by colocalization in at least one microglia population with eQTL H4 score (darker red is more significant). Results are filtered for protein coding genes only, full results are in Table S3.

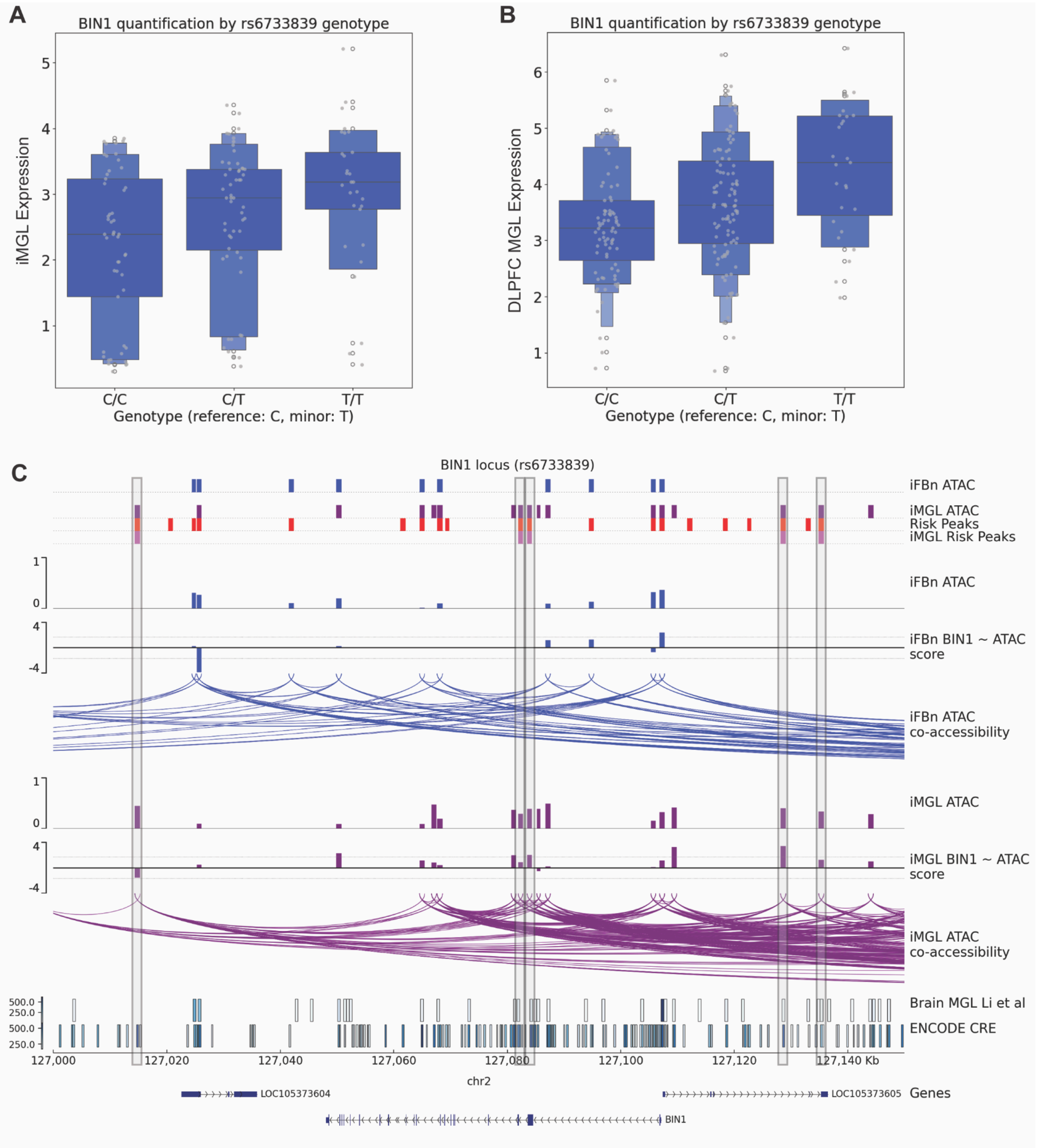


**Supplementary Figure 8:** BIN1 expression and locus characterization. A) Boxen/Scatter plot of BIN1 gene expression level by genotype in iMGL per allele dosage for the AD/LBD risk index variant rs6733839. B) BIN1 expression by genotype in single-nucleus microglia data from the DLPFC from the ROSMAP cohort. C) BIN1 locus tracks figure for co-accessibility of open chromatin peaks relative to cell-type and peaks containing associated risk variants. Blue arcs represent co-accessibility in iFBn and purple arcs show co-accessibility in iMGL. Shaded regions highlight co-accessibility in regions where peaks were selected for functional analysis.

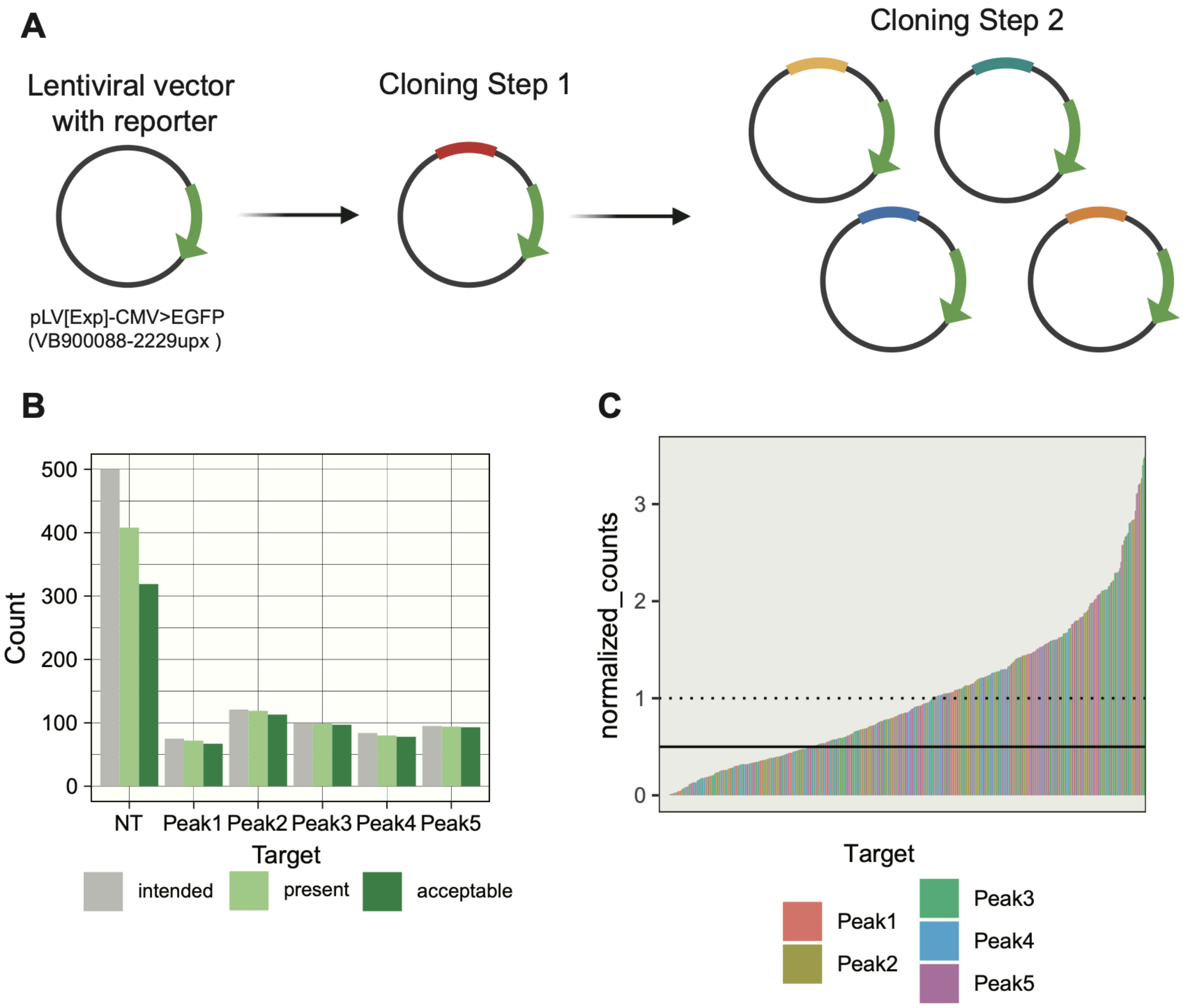


**Supplementary Figure 9:** Cloning strategy and QC. A) Guide machinery inserted into lentiviral plasmid with GFP reporter (pLV[Exp]-CMV>EGFP; VectorBuilder VB9000088-2229upx) in two steps. First, guide machinery with a holdover guide (scramble) inserted into the lentiviral vector (See Methods, Table S6). Second, pooled cloning removes the holdover guide and inserts guides of interest (Table S7). B) Distribution of pooled gRNAs for BIN1 peak-targeting experiment measured by amplicon sequencing of plasmid pool following cloning step 2 (See Methods, Table S6). Intended guides determined by gRNA sequences ordered for cloning (Table S7). Present guides determined by at least 1 read from amplicon sequencing aligning with 0% mismatch to intended guide sequence. Acceptable guides determined by at least 50% intended relative abundance in the pool (i.e. 1 guide in 10 guide pool = 10% theoretical abundance, “acceptable” if measured abundance > 5%). C) Measured relative abundance of peak-targeting guides in amplicon sequencing (dotted: theoretical abundance, solid: “acceptable” cutoff for relative abundance).

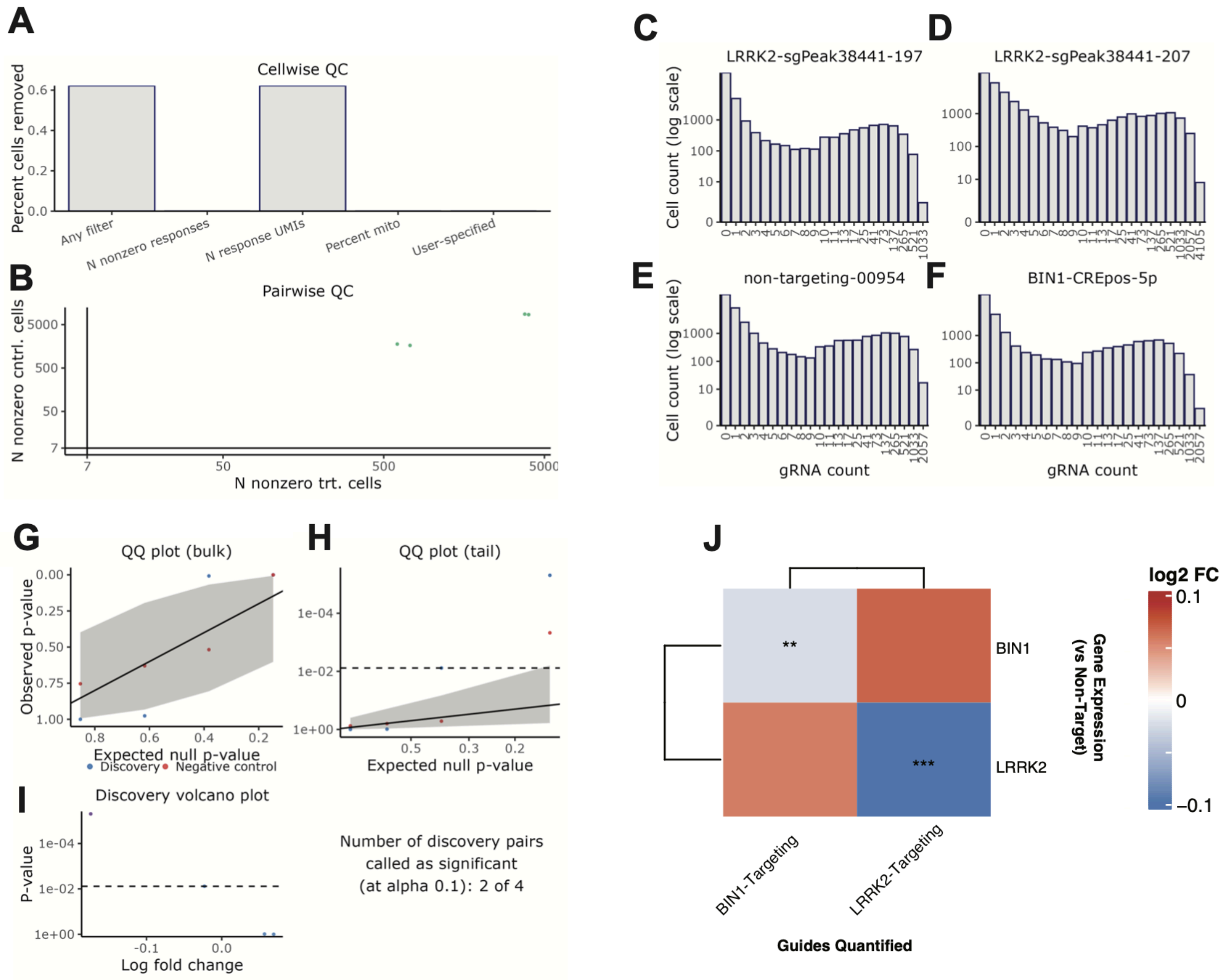


**Supplementary Figure 10:** LRRK2 and BIN1 promoter pilot. Target-feature correlation performed in SCEPTRE with "High-MOI" default parameters (See Methods). Cell-wise (A) and pairwise (B) quality control metrics reveal appropriate cutoffs for guides-in-cell and guides-per-target. (C-F) Representative distribution of guide counts per cell. (G-I) Target-feature association quality control metrics as provided by SCEPTRE. (J) Fold change in feature versus collective action of targeted guides (*** p_unadjusted < 0.001, ** p_unadjusted < 0.01).

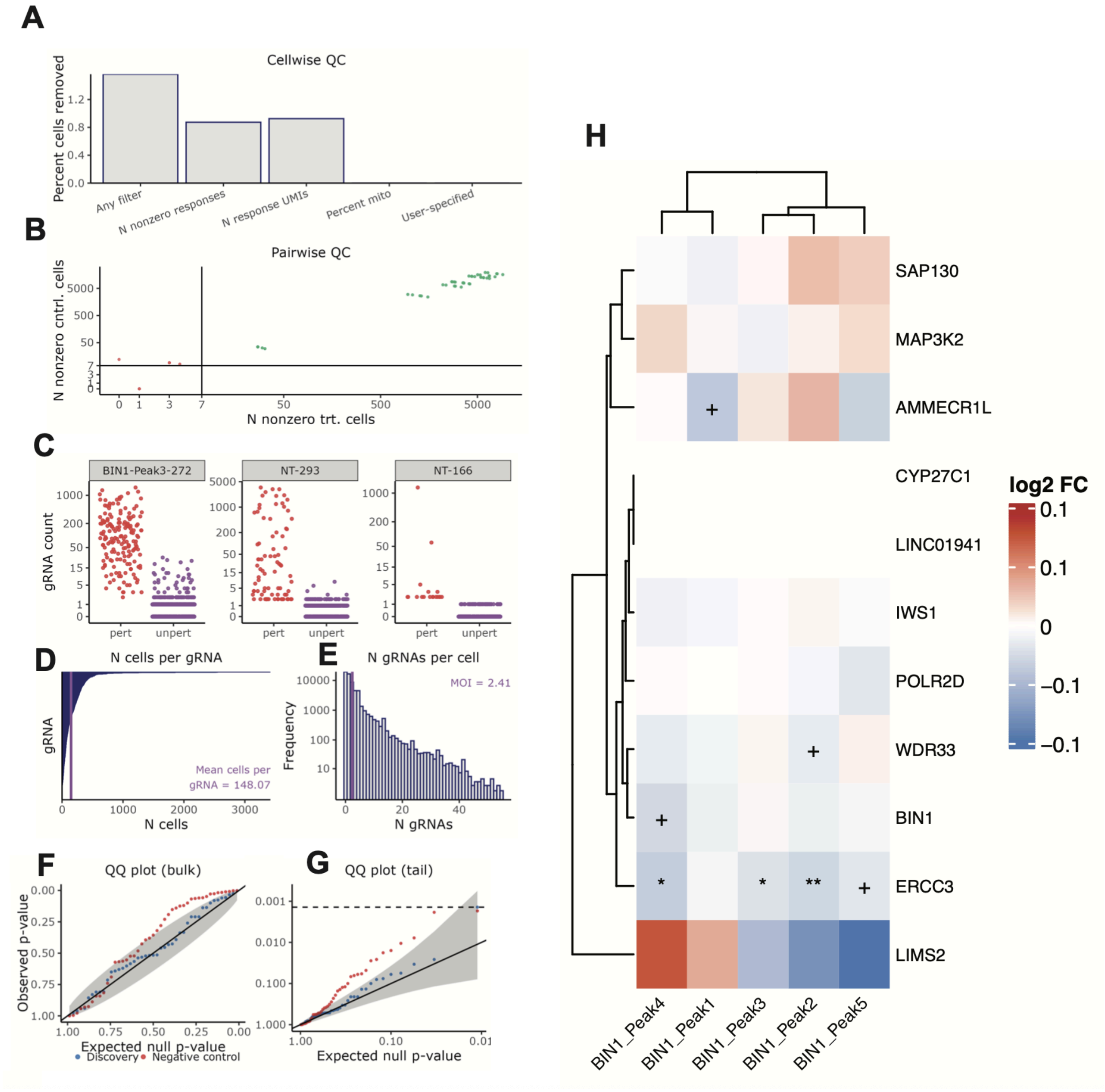


**Supplementary Figure 11:** SCEPTRE Analysis of candidate enhancer inhibition. Target-feature correlation performed in SCEPTRE with “High-MOI” default parameters (See Methods). Cell-wise (A) and pairwise (B) quality control metrics reveal appropriate cutoffs for guides-in-cell and guides-per-target. (C) Representative distribution of targeting versus non-targeting guide counts per cell. (D-G) Target-feature association quality control metrics as provided by SCEPTRE. (H) Fold change in feature versus collective action of targeted guides (** p_unadjusted < 0.01,* p_unadjusted < 0.05, + p_unadjusted < 0.1).

| Variable | Est. (95% Conf. Int.) | P-value | N |
|---|---|---|---|
| iMGL | -0.01(-0.04 to 0.02) | 0.576 | 133 |
| Kosoy meta-MGL | -0.09(-0.23 to 0.04) | 0.178 | 391 |
| Bryois MGL | -0.00(-0.13 to 0.13) | 0.993 | 192 |
| ROSMAP DLPFC MGL | 0.01(-0.04 to 0.06) | 0.606 | 206 |
| meta-MGL | -0.08(-0.25 to 0.09) | 0.342 | 730 |

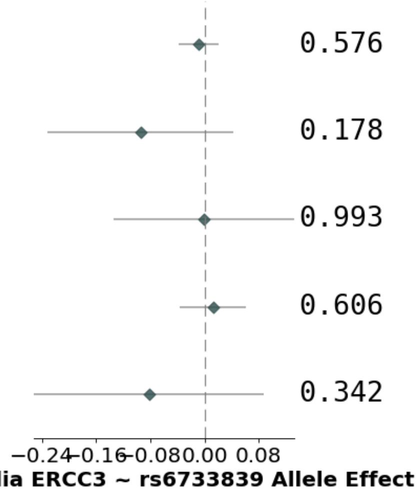


**Supplementary Figure 12:** caQTL correlated with *ERCC3* and BIN1 expression does not colocalize with genetic risk of AD. Forest plot in different MGL populations shows no colocalization of *ERCC3* with AD risk.
